# Consensus DNA Inhibits p53 Aggregation but Fails to Rescue Mutants

**DOI:** 10.1101/2025.07.24.665437

**Authors:** Matthew Steinsaltz, Gregory-Neal W. Gomes, Zachary A. Levine

## Abstract

The tumor suppressor p53 plays a crucial role in regulating gene expression under cellular stress. Somatic mutations to its DNA-binding domain are common in lethal cancers and lead to lost regulatory function and pathological hallmarks, such as misfolded p53 aggregates and amyloid-like fibrils. Despite decades of these observations, the molecular determinants driving p53 aggregation and its role in cancer incidence, progression, and lethality remain poorly defined. Identifying these determinants, however, is critical for developing therapeutic interventions that modulate protein solubility in cancer and understanding disparities in cancer outcomes. Therefore, we investigated whether consensus DNA sequences that stabilize wild-type p53 are sufficient to regulate the aggregation of oncogenic p53 mutants with reduced binding affinities. Using a combination of probe-based and label-free spectroscopic and microscopic techniques, we examined how consensus (p21, Bax) and non-consensus (p21-scramble, Poly-GC) DNA oligonucleotides regulate the aggregation of wild-type p53 in comparison to three cancer-associated mutants (R248Q, R273H, and R175H). We find that equimolar p21 consensus sequences significantly inhibits wild-type p53 aggregation, while other DNA sequences do not. In contrast, oncogenic p53 mutants evaded DNA regulation of protein aggregation, and some oligonucleotides even enhanced aggregation at low concentrations, which suggests concentration-dependent DNA-p53 interactions. These findings emphasize that DNA response elements are sufficient for regulating wild-type p53 aggregation in solution. However, key somatic mutations in cancer promote aggregation at the expense of DNA-binding, directly leading to the loss of p53 solubility through biochemical interactions. Taken together, these observations suggest that potent regulators of p53 aggregation should aim to restore affinity between oncogenic p53 mutants and regulatory DNA to minimize the pathological hallmarks of lethal tumors.

**Significance Statement:** Our research investigates whether DNA sequences that bind tumor suppressor protein p53 regulate its aggregation behavior in wild-type and somatically mutated forms linked to cancer. While specific DNA sequences regulate p53 aggregation, protein mutations that reduce affinity for consensus DNA or destabilize p53’s conformation result in persistent, likely dysfunctional aggregates, even in the presence of regulatory nucleotides. Unsurprisingly, p53 and it’s mutants exhibit increased aggregation when there is substantially more protein to DNA, suggesting insufficient regulatory DNA or excess unregulated p53 may inevitably lead to pathological aggregation. These findings clarify our understanding of p53’s molecular behavior and suggest that cancer-linked somatic mutations enable p53 to evade DNA’s regulatory effects and subsequently aggregate, leading to downstream losses in function.

## Introduction

As a transcription factor and tumor suppressor, p53 functions as a homotetramer (1, 2) to regulate genes that are critical for cell cycle control, DNA repair, and apoptosis through sequence-specific DNA binding (3, 4). However, these functions are inactivated by mutations in approximately 30-50% of all cancers (5, 6), with frequencies exceeding 90% in various lethal cancers such as ovarian carcinomas (7, 8). Analysis of data generated from The Cancer Genome Atlas (TCGA) Research Network (https://www.cancer.gov/tcga) through cBioPortal (9–11) reveals these mutations are primarily missense alterations that cluster within the DNA-binding domain (DBD) of p53 (Fig. 1A,B; PDB: 1TUP), with the most frequent occurrences at residues R175, G245, R248, R249, R273, and R282 (Fig. 1C). Mutations to these DNA binding sites can be classified as either contact mutants (R248Q and R273H), which directly reduce DNA-binding affinity, or conformational mutants (R175H), which structurally destabilize p53 through impaired zinc coordination (12–15). These p53 mutations often result in a spectrum of functional consequences, ranging from partial deficiencies and dominant-negative effects to a complete loss of tumor-suppressive function (loss-of-function) (16, 17) as well as the acquisition of new pro-oncogenic characteristics (gain-of-function) (18–20).

**Figure 1.**
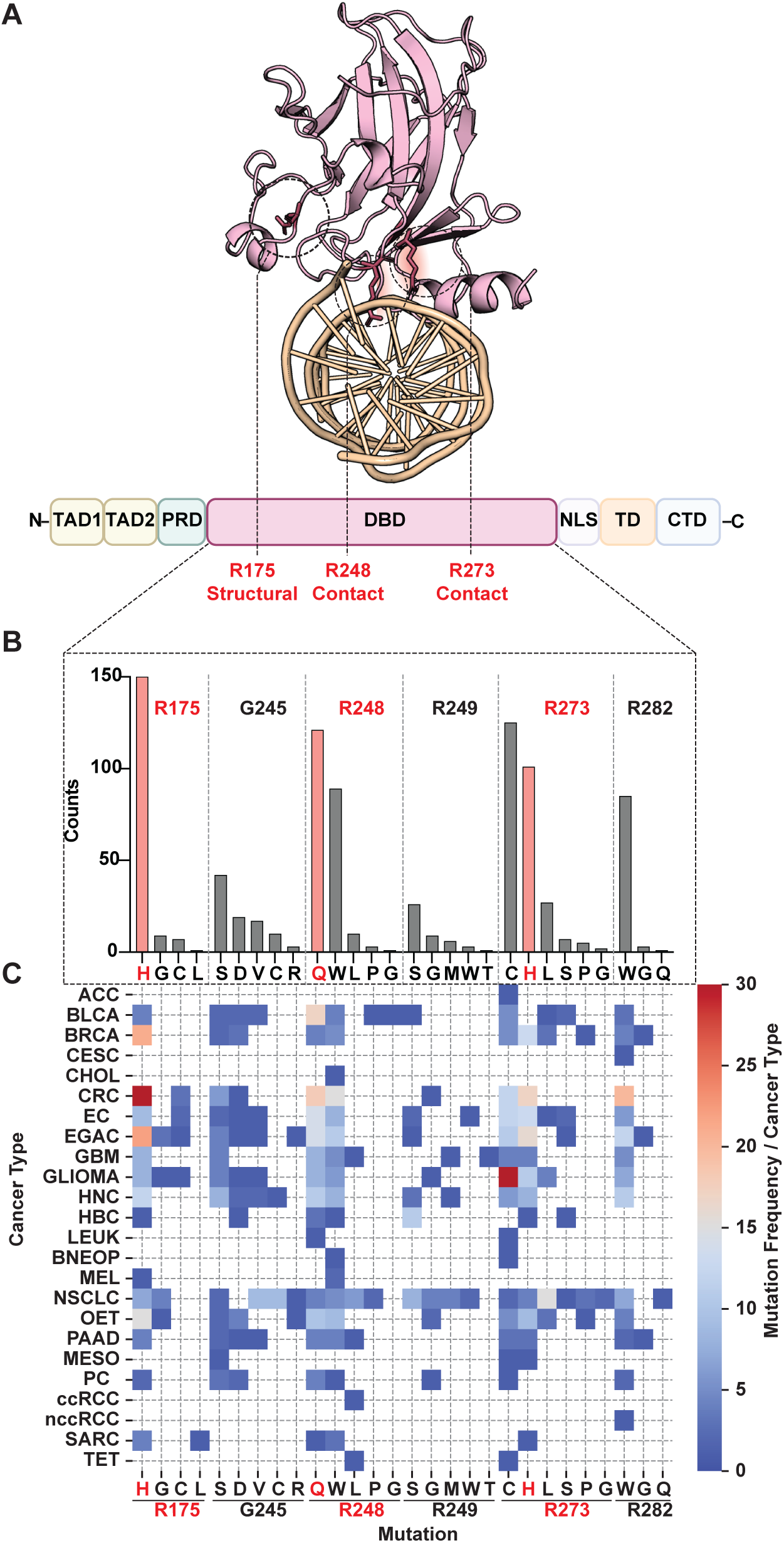
Mutations within the p53 DBD are highly prevalent across cancers and frequently occur at Arg residues critical for structure and function. A) Structure of monomeric p53 DBD (pink) interacting with DNA (brown) from PDB: 1TUP. Residues R175, R248, and R273 mutated in the DBDs used in this study are colored and highlighted in red. Domains in p53 schematic are as follows: Transactivation domains (TAD1-2); Proline-rich domain (PRD); DNA binding domain (DBD); Nuclear localization signal (NLS); Tetramerization domain (TD); C-terminal regulatory domain (CTD). B) Counts of somatic missense mutations at p53 hotspot residues from cBioPortals TCGA Pan Cancer Atlas. C) Heatmap of missense mutations across cancer types from TCGA Pan Cancer Atlas. Cancer acronyms are listed in table 1.

Although aggressive cancers often contain p53 missense mutations, the mechanisms underlying their oncogenic influence and context-sensitive impact on disease etiology and progression remain incompletely understood (17, 21, 22). Therefore, while intracellular levels of p53 are tightly and dynamically regulated according to cell state and stress (23), p53’s intrinsically low thermodynamic stability—further decreased by mutation—means that both WT and, particularly, specific mutant p53 proteins can readily misfold (15, 24–27). If not properly degraded, they can interact and form irreversible amyloid-like aggregates by exposing hydrophobic segments capable of forming steric zippers between multiple aggregation-prone interfaces (28–31). Biologically, p53 aggregates have been identified in multiple lethal tumor samples (32–36) and linked to poor prognoses (33). Moreover, aggregation can lead to a prion-like inactivation mechanism that traps and neutralizes other members of the tumor suppressor family, such as p63, p73, as well as tau (37–41), suggesting that the promiscuity of p53 aggregates may function as a tunable yet druggable driver of cancer lethality and neurodegeneration. Additionally, recent studies also indicate that p53 monomers can undergo liquid-liquid phase separation, forming reversible condensates that can transition into irreversible aggregates influenced by salt concentration, temperature, pH, post-translational modifications, and nucleic acid interactions (42–45).

Current therapeutic strategies to mitigate p53 aggregation largely focus on stabilizing its native conformation, inhibiting aggregation-prone sequences, or attempting to restore wild-type function in mutants using small molecules and peptides (40, 46–50). Within this context, nucleic acids, and specifically DNA, are a powerful, yet underexplored, modulator of p53 aggregation. Early work by Bullock et al. demonstrated that consensus double-stranded DNA could significantly increase the melting temperature of p53’s DNA-binding domain (26). Subsequent studies further showed that both the DBD and full-length p53 could be rescued from misfolding through specific interactions with various DNA response elements (51). Despite these foundational observations, a critical knowledge gap remains: it is unclear whether common cancer-associated missense mutations in p53 allow the protein to biochemically evade DNA’s regulatory influence on its stability and proclivity towards aggregation. Addressing this necessitates a systematic evaluation of how diverse and concentrated DNA response elements impact the aggregation propensity of both wild-type and mutant p53 (52). A deeper understanding of these sequence-dependent effects is crucial as p53 orchestrates stress-dependent gene regulation via response elements that exhibit differential binding affinities (53, 54). For example the p21 response element, vital for cell cycle control, binds p53 with higher affinity than the Bax response element involved in apoptosis (55). Furthermore, while factors like DNA flexibility (56) and accessibility (57, 58) have also emerged as potentially orthogonal regulators of the p53 response, differences in their native binding affinities will likely influence p53’s behavior in the cell, further motivating the need to compare interactions between diverse p53 mutants and DNA response elements. Moreover, the precise mechanisms by which mutant p53 evades endogenous regulation, leading to its elevated levels in tumor cells—a key factor in many of its oncogenic effects—remain unresolved (57–60). This overall ambiguity hinders the development of tailored therapeutic strategies to alleviate the burden of mutant p53, as evidenced by the current lack of FDA-approved drugs targeting the mutant form of the protein (16, 61).

To address these unmet needs, we utilized a combination of biophysical and biochemical tools to probe the aggregation of multiple p53 DBD mutants in the presence of diverse DNA response elements and test the hypothesis that mutations to p53 confer resistance to DNA regulatory elements through a concentration-dependent mechanism. Our findings support and expand on earlier reports that specific DNA oligos (e.g., p21-RE) can reduce aggregation of WT p53 DBD in a sequence-dependent manner, but only at equimolar ratios. Interestingly, lower DNA concentrations enhanced mutant (R248Q and R273H) aggregation, consistent with protein-RNA observations where low RNA concentrations promoted aggregation while high concentrations reduced aggregation (62). Through a combination of orthogonal spectroscopic techniques, we demonstrate that while p21-RE effectively reduces WT p53 DBD aggregation, no combination of DNA consistently regulates mutant p53 aggregation. Finally, a systematic comparison of our techniques illustrates that changes in DNA concentration or sequence have less of an effect on p53 aggregation than the effect of p53 mutations alone. These observations indicate that while DNA is sufficient to reduce WT p53 aggregation (likely through the stabilization of native states at equimolar concentrations), the inherent structural instability of mutant p53 renders it resistant to endogenous DNA regulation. This suggests that exogenous therapeutic strategies should aim to increase DNA response element binding to best restore p53’s loss in native function and minimize the hallmarks of cancer and aging.

## Results

### Sequence-specific DNA regulation of WT p53 DBD aggregation

To investigate DNA’s regulatory effects on p53 aggregation, we first established a robust experimental framework through careful quality control steps. After validating proper protein folding and WT-DNA binding, we established complementary techniques to monitor aggregation in the presence of DNA. These methodological refinements enabled us to systematically examine how DNA sequence specificity and concentration influence the aggregation behavior of the p53 DBD. Initial thermal denaturation studies confirmed proper folding of purified WT p53 DBD, with a melting temperature (Tm) of 42.57°C (Fig. S1), consistent with previously reported values (26, 63). We established experimental conditions using equimolar (1:1) p53:DNA as our highest intervention ratio at 5μM, and tested two lower DNA concentrations (9:1 and 454:1 protein:DNA) to examine concentration-dependent effects (45, 52). Gel shift assays demonstrated that p21-RE maintained consistent binding across all ratios, while Bax, p21sc (scrambled p21), and PolyGC sequences showed minimal binding (Fig S2A). The expected binding hierarchy (p21-RE > Bax > p21-scramble > PolyGC) was evident initially (Fig S2B), but only p21-RE maintained binding over extended periods (24 hrs to one month) (Fig S2C). Notably, p21-RE binding persisted in supernatant fractions of aggregated samples, suggesting DNA-bound p53 DBD likely becomes unavailable for aggregation. While lower affinity sequences poorly retained p53 DBD, they may still influence aggregation dynamics during early timepoints.

With our experimental conditions and DNA binding confirmed, we then proceeded to probe p53’s aggregation kinetics with Thioflavin-T (ThT), a dye with greatly enhanced fluorescence when bound to β-sheet rich structures such as amyloid fibrils (64). While ThT effectively monitored p53 DBD aggregation in isolation, its positive charge and known DNA intercalation properties (65–67) complicated measurements in the presence of DNA. Indeed, control experiments revealed significant ThT fluorescence quenching by DNA alone, precluding its use in our DNA-dependent aggregation studies. We therefore employed an alternative commercial dye from the luminescent conjugated oligothiophene family, AmyTracker630 (AT630), which are characteristically negatively charged at physiological pH and designed to bind positively charged grooves within amyloid fibrils and prefibrillar aggregates (68–70). It is likely due to its predicted negative charge that AT630 avoids interference with DNA while producing robust fluorescence signals from protein aggregates both in the presence and absence of DNA, with little response to DNA alone (Fig. S3B). Signal-to-noise analysis showed that DNA produces only a ∼two-fold increase in fluorescence over background (dye alone), while protein aggregates produce an approximately 200-fold increase after 24 hours, indicating substantially higher AT630 sensitivity to protein aggregates compared to DNA. In subsequent kinetic studies, substoichiometric dye concentrations were employed to minimize potential dye-mediated effects on the aggregation process (68, 71) while maintaining robust signal detection due to AT630’s high sensitivity.

The addition of equimolar p21-RE considerably slowed p53 DBD aggregation, extending the mean aggregation half-time from 166 ± 44 (protein alone) to 490 ± 60 minutes (mean ± SD) (Fig. 2A). Other DNA sequences showed more modest effects, with Bax and p21-scramble yielding half-times of 234 ± 29 and 234 ± 36 minutes respectively, and PolyGC at 189 ± 40 minutes. The effect of DNA varied with protein:DNA ratios: while equimolar conditions slowed aggregation and reduced final aggregate amounts (Fig S4A), low p21-RE concentrations (protein:DNA = 454:1) accelerated aggregation to a half-time of 108 ± 13 minutes. This acceleration at high protein:DNA ratios was observed for p21-RE, p21sc, and PolyGC sequences, though notably without affecting the final aggregate amount compared to DNA-free samples (Fig S4B). Complete statistical analyses of AT630 measurements are provided in Table S1. To orthogonally confirm these findings and directly visualize aggregate morphology, we employed fluorescence confocal microscopy combined with transillumination on equimolar p21-RE, which yielded the most prominent reduction in aggregation (Fig. 2C). While transillumination alone revealed distinct aggregate structures, AT630 fluorescence specifically labeled p53 aggregates without binding to DNA, confirming their amyloid nature. Quantification using the SINAP deep learning model (InCarta software) (Fig. S5) across three independent wells demonstrated higher fibril confluence (surface area coverage) for WT DBD alone compared to samples with equimolar p21-RE (Fig. 2D), corroborating our kinetic spectroscopic findings.

**Figure 2.**
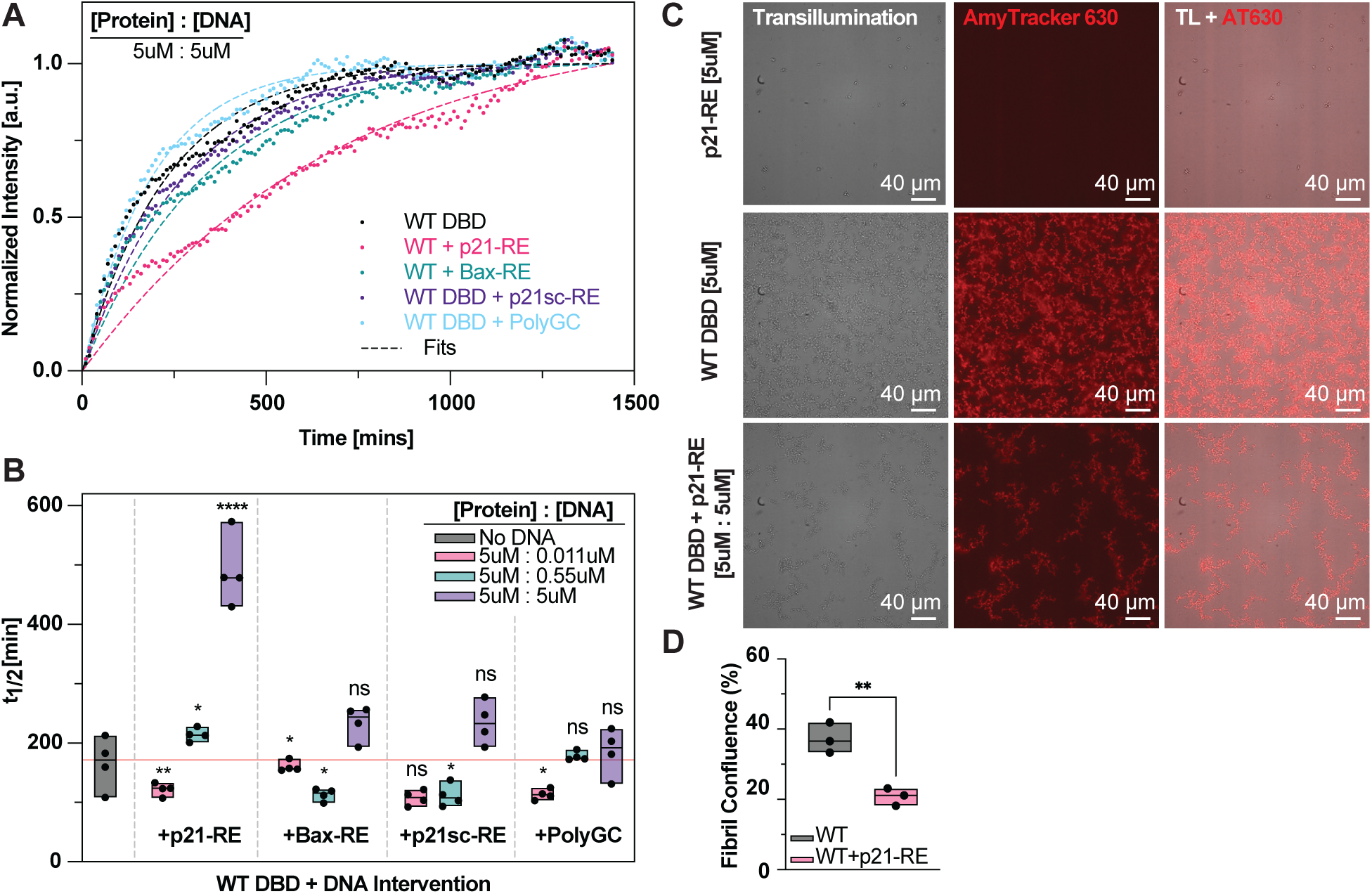
WT p53 DBD aggregation is DNA sequence- and concentration-dependent as measured by amyloid-specific dye. A) Representative time course curves and fits of normalized AmyTracker630 intensity for WT p53 DBD over 24 hours with and without consensus DNA at equimolar protein to DNA concentration (5 : 5 μM) (n = 4). B) The t-half values for WT p53 DBD were obtained from independent one-phase association fits to each replicate. All sequences for each concentration ratio were analyzed using a one-way ANOVA followed by a post hoc Dunnett’s test with the untreated sample as the control. If no symbols are displayed, the sample failed to reject the ANOVA. C) WT p53 DBD aggregation decreases in the presence of equimolar p21-RE via imaged-based analysis. Transillumination and AmyTracker630 fluorescence of p53 aggregates at 22hrs with and without p21-RE captured via confocal microscopy (n = 3). D) Fibril confluence (surface area covered by fibrils) calculated with the AmyTracker630 channel using the SINAP deep-learning model within the IN Carta software from Molecular Devices, followed by a student t-test. P-values were mapped to significance symbols as follows: ns ≥ 0.05 ; * : P ≤ 0.05 ; ** : P ≤ 0.01 ; *** : P ≤ 0.001 ; **** : P ≤ 0.000

Next, to increase the confidence of our observations and minimize artifacts associated with dyes or fluorophores, we also employed intrinsic fluorescence measurements of the single tryptophan residue in p53 DBD (Fig. 3D), which served as an orthogonal probe for conformational changes during aggregation. Among all tested sequences, equimolar p21-RE induced the largest reduction in intrinsic fluorescence, decreasing the signal from 2976 ± 186 a.u. for DBD alone to 1642 ± 42 au., compared to more modest reductions seen with Bax, p21-scramble, and Poly(GC) sequences (Fig. 3A). This pronounced decrease in fluorescence suggests DNA-mediated stabilization of native or near-native conformations. Initial fluorescence measurements were consistent across all conditions (∼500 a.u.; Fig. 3B). After 24 hours at 37°C, all DNA-containing samples showed increased intrinsic fluorescence compared to WT DBD alone. This increase followed two distinct patterns: a concentration-dependent effect of DNA (5 μM > 0.55 μM > 0.011 μM) and a sequence-specific trend (p21 < Bax ≲ p21sc < PolyGC) relative to WT DBD alone. Therefore, the minimal changes in tryptophan fluorescence across all conditions at initial time points indicate that DNA binding alone does not significantly alter p53 DBD conformation. However, the substantial increase in fluorescence after 24 hours suggests that progressive protein misfolding, rather than DNA binding, drives the major conformational changes in p53 DBD structure. Results for all trp fluorescence statistical analyses can be found in Table S2.

**Figure 3.**
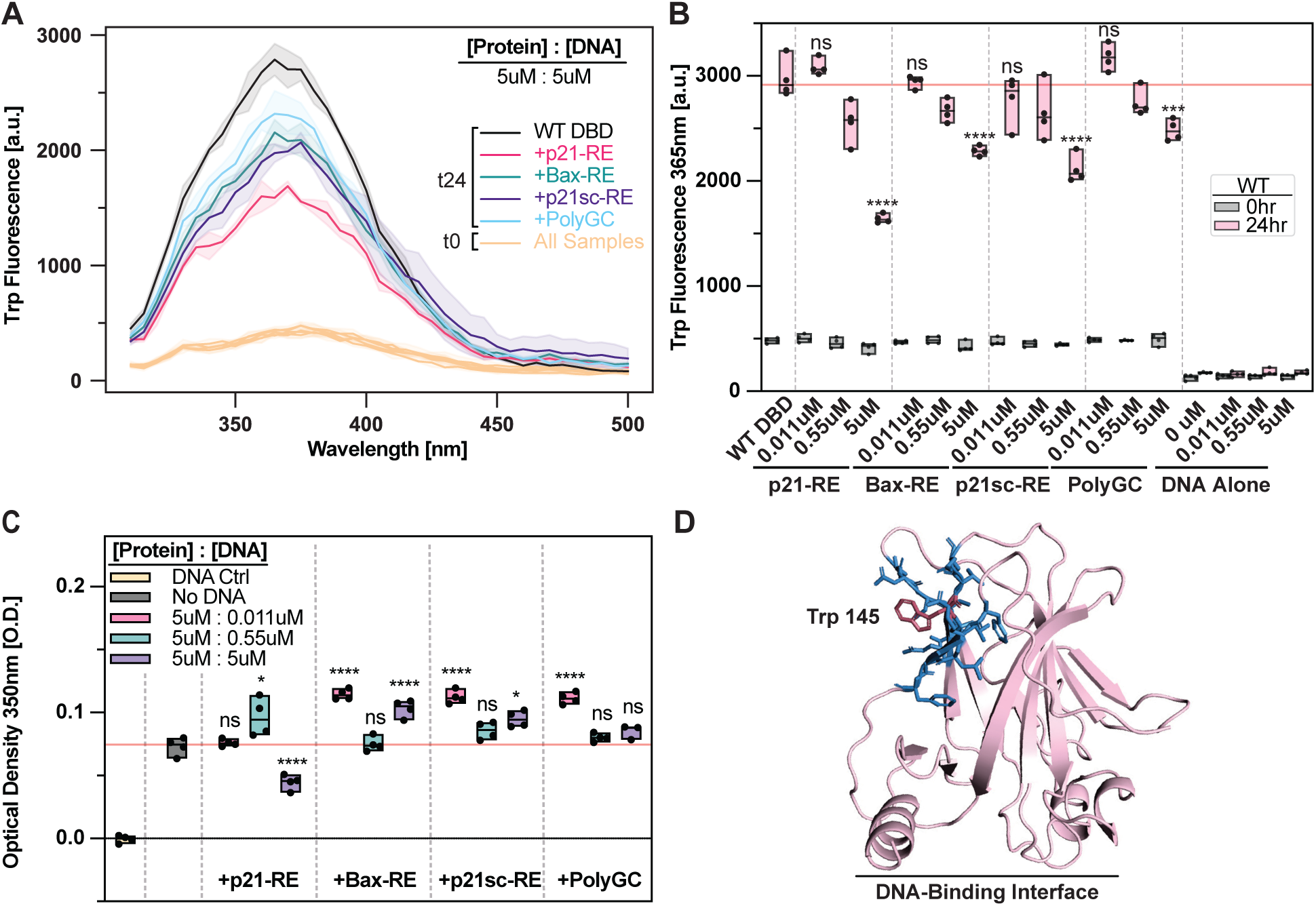
Label-free spectroscopic methods support DNA sequence- and concentration-dependent modulation of WT p53 DBD aggregation. A) Emission Scan for Internal Trp Fluorescence (n = 4). B) Boxplots of emission values at 365nm. Internal tryptophan fluorescence was measured in quadruplicate before aggregation (0 hours) and after 24 hours of incubation at 37°C to initiate aggregation. Samples were excited at 298nm and scanned from 310 to 500nm. C) Optical density of homogenized sample after aggregating conditions at 350nm (n = 4). D) Monomeric p53 DBD (PDB: 1TUP) displayed in pink, highlighting the single Trp residue relatively solvent exposed in red, with surrounding residues in blue. All sequences for each concentration ratio were analyzed using a one-way ANOVA followed by a post hoc Dunnett’s test with untreated sample as the control. If no symbols are displayed, the sample failed to reject the ANOVA. Dunnett’s adjusted P-values were mapped to significance symbols as follows: ns ≥ 0.05 ; * : P ≤ 0.05 ; ** : P ≤ 0.01 ; *** : P ≤ 0.001 ; **** : P ≤ 0.000

To further confirm these observations using an additional label-free method, we analyzed changes in optical density, a technique routinely employed in kinetic analyses that also serves as an informative endpoint measurement (72). Consistent with tryptophan fluorescence, equimolar p21-RE uniquely reduced optical density compared to DBD alone (Fig. 3C). However, higher amounts of p53 resulted in increased optical densities, suggesting the formation of more numerous or larger aggregates. While this initially appears to contrast the decreased AT630 fluorescence at similar ratios, this discrepancy can likely be explained by the formation of non-fibrillar aggregates that lack the specific fibrillar features necessary for AT630 binding, yet still contribute to nonspecific light scattering measured by optical density. Results for all optical density statistical analyses can be found in Table S3.

### p53 DBD mutants resist DNA-mediated stabilization

Building on our characterization of WT p53 DBD aggregation, we next examined three p53 DBD mutants prevalent in cancer. While WT p53 DBD shows reduced aggregation in the presence of consensus DNA such as p21-RE, the response of cancer-associated mutants was previously unclear. In particular, we wished to test whether contact mutants (R248Q and R273H) and conformational mutants (R175H) would escape DNA-dependent regulation. Given their reduced binding affinities, we also wanted to test the extent to which DNA-binding vs. p53 misfolding contributes to protein aggregation and whether DNA regulation can overcome one or both of these changes. We first characterized the DNA binding affinity and thermal stability of each of the p53 mutant-DNA pairs. According to gel shift assays, minimal binding was observed before or after aggregation with the exception of R248Q (Fig. S6). Intriguingly, thermal melt analysis (Fig. S1) revealed that R248Q was slightly more thermally stable than WT, which may explain its somewhat reduced aggregation propensity – an atypical finding that has been previously reported and attributed to differences in buffer conditions between studies (13). R273H showed only marginally reduced stability compared to WT (27), with both variants displaying a higher Tm than previously demonstrated by Khadiullina et al. (63), whereas R175H exhibited a considerably lower thermal stability.

Next, we characterized the aggregation kinetics using AT630. Each p53 mutant displayed distinct kinetic profiles, with aggregation half-times following the order R175H < R273H < R248Q (21 ± 1.7, 86 ± 22, and 425 ± 84 minutes, respectively) (Fig. S7). The introduction of DNA largely did not extend these half-times but rather decreased the half-times in some, but not all, high protein-to-DNA ratios for R248Q and R273H (Fig 4A,B,C), thereby accelerating aggregation. The relative rates of aggregation matched the final aggregate intensities among the mutants: R248Q showed both the slowest aggregation and lowest final intensity, while R273H and R175H displayed faster kinetics and higher final intensity values (Fig S8). DNA most strongly influenced R273H’s aggregation behavior, promoting the formation of larger or more abundant aggregates at high and intermediate protein:DNA ratios, despite only modestly accelerating the aggregation rate. In contrast, WT p53 showed minimal changes in final aggregate size or abundance even when DNA slightly accelerated its aggregation at low concentrations. While high protein:DNA ratios led R248Q and R273H to form somewhat larger or more numerous aggregates more quickly, R175H’s aggregation remained largely unaffected by DNA.

**Figure 4.**
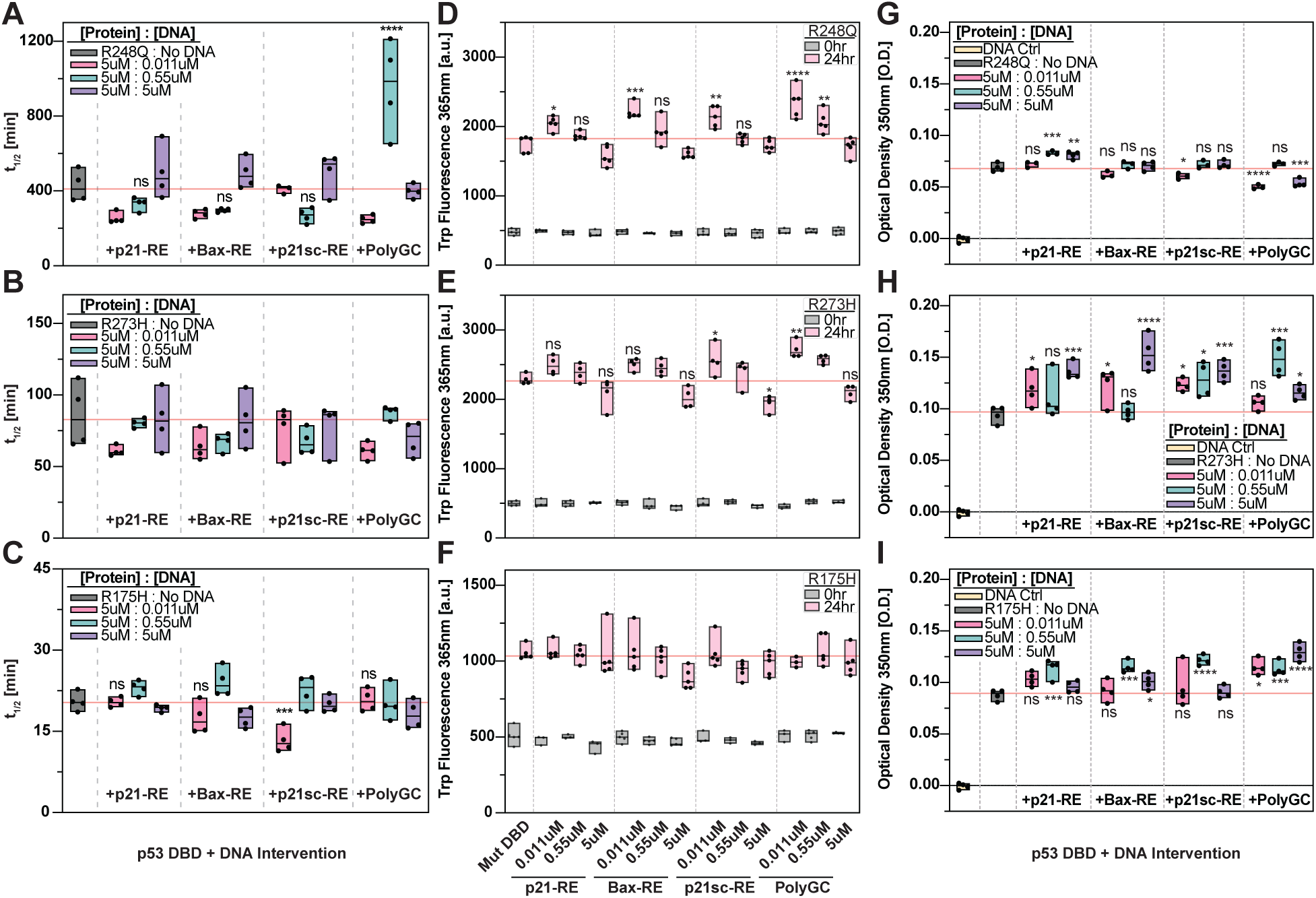
Mutant p53 DBDs exhibit diminished DNA sequence sensitivity and modest enhanced aggregation at low DNA concentrations. A) The t-half values for R248Q, B) R273H, and C) R175H p53 DBDs obtained from independent one-phase association fits to each replicate in each sample (n = 4). Internal tryptophan fluorescence at 365nm for D) R248Q, E) R273H, and F) R175H before and after aggregation (n = 3-5). G) Optical density of homogenized sample before and after aggregating conditions at 350nm for R248Q, H) R273H, and I) R175H (n = 3-4). All sequences for at each concentration ratio for each protein were analyzed using a one-way ANOVA followed by a post hoc Dunnett’s test with untreated sample as the control. If no symbols are displayed, the sample failed to reject the ANOVA. Dunnett’s adjusted P-values were mapped to significance symbols as follows: ns ≥ 0.05 ; * : P ≤ 0.05 ; ** : P ≤ 0.01 ; *** : P ≤ 0.001 ; **** : P ≤ 0.000

Beyond kinetic profiles, internal tryptophan fluorescence provided further insights into DNA’s influence on mutant p53 aggregation. For R248Q and R273H, we observed a concentration-dependent, but not sequence-dependent, trend in DNA’s ability to modulate aggregation, similar to WT p53 (Fig 4D,E,F and Fig S9). However, their response to low DNA concentrations noticeably differed. While WT samples at low DNA concentrations yielded signals close to the protein-only baseline, R248Q and R273H showed significant signal elevations. PolyGC, for example, produced the highest signals (2350 ± 223 a.u. for R248Q and 2718 ± 129 a.u. for R273H) compared to their respective protein-only values (1779 ± 112 a.u. and 2289 ± 76 a.u.). This initial elevation was followed by a concentration-dependent decrease in signal for each oligonucleotide, mirroring WT behavior. In stark contrast, R175H showed minimal response to DNA addition; its signals across all concentrations remained tightly clustered around its protein-only values, indicating no consistent concentration-dependent trend. These distinct fluorescence patterns suggest that while DNA can influence some p53 mutants, it largely fails to exert the same regulatory, aggregation-inhibiting effect observed with WT p53, particularly on structural mutant R175H. Adding another layer to these observations, optical density measurements also highlighted key differences between wild-type and mutant responses, generally showing inconsistent and heterogeneous responses to DNA across conditions (Fig 4G,H,I). The lack of clear patterns in optical density likely reflects the combination of weaker DNA-mediated effects on mutant aggregation and the limited sensitivity of these bulk measurements. Furthermore, across all three assays, we consistently found no sequence-dependent reduction in aggregation for mutant p53 DBDs, directly contrasting our clear observations with WT p53 DBD. Interestingly, high protein:DNA ratios (454:1) did appear to slightly, yet inconsistently, enhance aggregation for both WT and mutant samples.

### Mutant-specific patterns dominate over DNA-dependent effects

To comprehensively analyze p53 DBD behavior across different mutations and conditions, we integrated data from three complementary techniques: AmyTracker630 kinetics, internal tryptophan fluorescence, and optical density measurements. Each technique provides distinct insights: AT630 reports on aggregate formation and structure through dye binding, tryptophan fluorescence reveals local protein conformational changes, and optical density measures overall aggregate size and quantity. When examining the multi-dimensional dataset, distinctive distribution patterns emerged when median values for each condition were grouped by DBD (Fig 5A-C). These patterns reflect the complementary nature of our measurements: AT630 data clusters reveal similarities in amyloid-like structure formation, optical density groupings indicate shared aggregate size distributions, and tryptophan fluorescence patterns suggest common local structural changes. Most strikingly, the WT + equimolar p21-RE condition (gold circle and arrow, Fig 5) consistently appeared separated from other WT conditions and positioned closer to R248Q measurements across all three measurement spaces. This consistent positioning across fundamentally different measurement techniques provides strong evidence for p21-RE’s unique ability to modulate WT p53 DBD aggregation.

**Figure 5.**
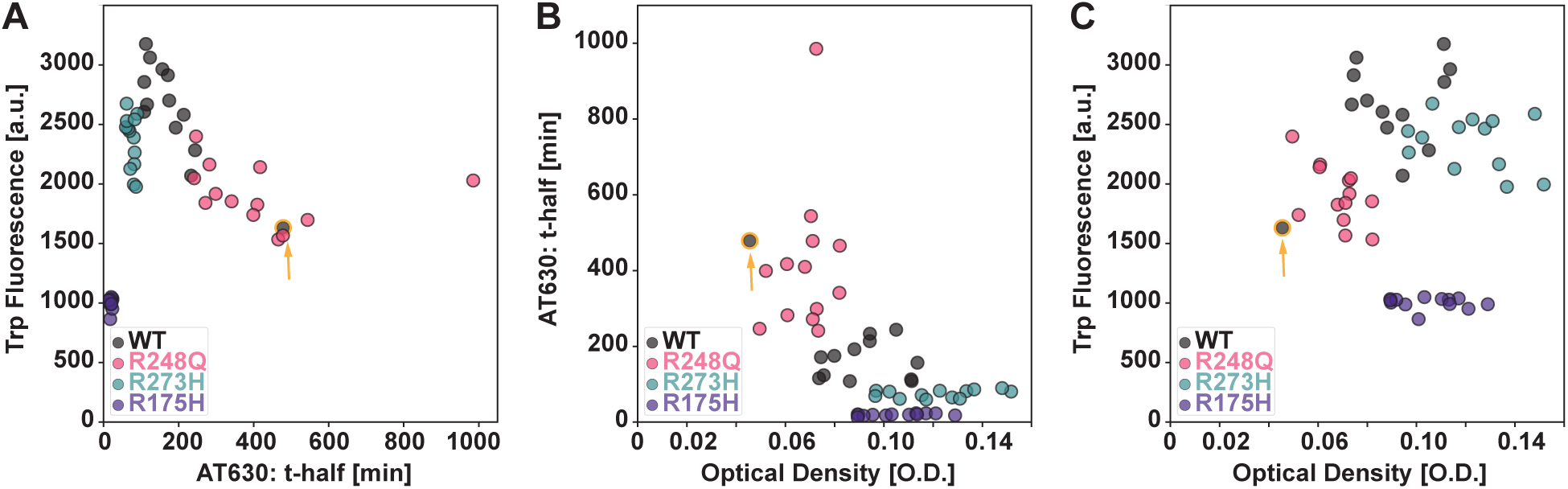
WT and mutant p53 DBDs (R248Q, R273H, R175H) show distinct patterns across multiple aggregation measurements. A) Inter-method comparison for AT630 vs. Trp Fluorescence; B) Optical Density vs. AT630; and C) Optical Density vs. Trp Fluorescence. Each point represents the median of each sample’s replicates and is grouped by DBD. Equimolar WT + p21-RE is highlighted in gold and indicated by a gold arrow.

Crucially, further analysis of these clustering patterns revealed that mutation-specific characteristics largely dictated aggregation behavior across techniques, often overshadowing DNA-dependent effects. In AT630 kinetics, R175H and R273H measurements clustered together but separated from WT and R248Q, which may suggest similar aggregation pathways for these two mutants. Tryptophan fluorescence data revealed distinct groupings, with R175H measurements clustering with R248Q, suggesting similar local structural changes despite differing global aggregation patterns. Optical density measurements revealed yet another pattern, with R248Q clustering closer to WT, while R175H and R273H showed substantial overlap, suggesting similar aggregate size distributions for these mutants. These technique-specific clustering patterns, largely driven by the inherent properties of each mutation, consistently suggest that distinct p53 mutations lead to different aggregation behaviors or propensities. This observation highlights how mutation-specific behaviors enable p53 to evade the regulatory effects of DNA on aggregation, underscoring the need for tailored therapeutic approaches that address the unique pathology of each mutant.

## Discussion

### DNA-Mediated Modulation of WT p53 DBD Aggregation

WT p53 DBD aggregation showed clear sequence- and concentration-dependent responses to DNA interventions. At equimolar concentrations, p21-RE specifically reduced aggregation across multiple experimental measures, consistent with previous observations of DNA-mediated stabilization (51). However, lower DNA concentrations produced slightly increased aggregation, following patterns similar to those observed in protein-RNA interactions where concentration ratios dictate assembly versus dissolution of protein condensates (52, 62). This biphasic response could be explained by several potential mechanisms: while high DNA concentrations promote sequestration-based inhibition, low DNA concentrations may enhance aggregation through partial DNA binding that blocks certain aggregation interfaces while exposing others (similar to multisite aggregation for p53 peptide interactions (31), or by creating local nucleation points that concentrate proteins. Alternatively, sparse DNA molecules could alter the local electrostatic environment, causing proteins to reorient in ways that expose interaction-prone regions, thereby facilitating protein-protein contacts rather than protein-DNA binding. Intriguingly, while mutant p53 DBD variants showed inconsistent responses overall, some demonstrated increased aggregation at high DNA concentrations, suggesting that even impaired DNA binding might be sufficient to trigger nucleation-based aggregation without achieving the stabilizing effects seen with WT protein (Fig. 6).

**Figure 6:**
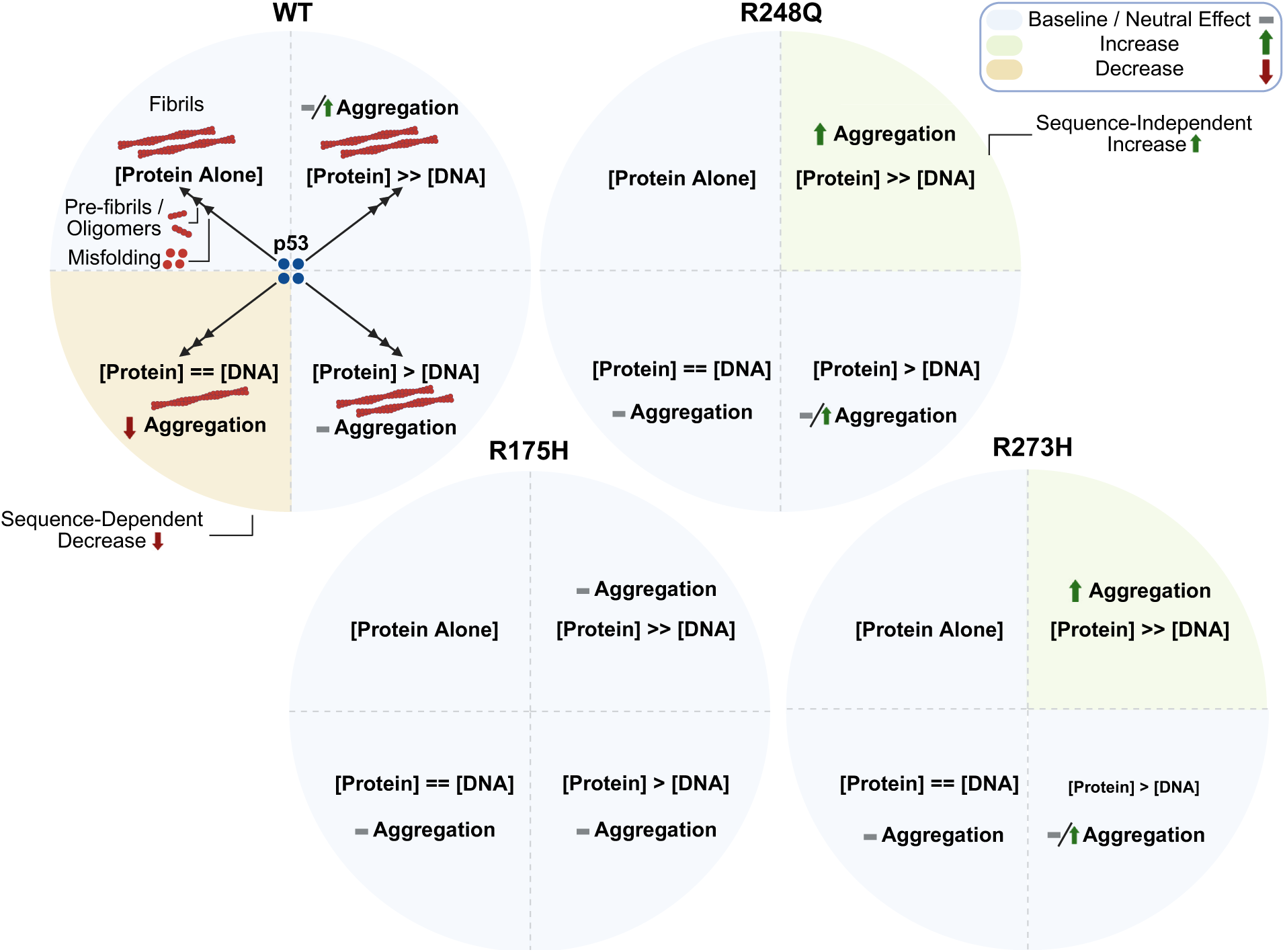
Differential effects of DNA on WT and mutant p53 DBD aggregation pathways. DNA modulates p53 DBD aggregation based on both protein variant and DNA concentration, with sequence specificity uniquely affecting WT p53 at equimolar ratios. Three key protein:DNA ratios were examined: high ([Protein] >> [DNA], 454:1), intermediate ([Protein] > [DNA], 9:1), and equimolar ([Protein] == [DNA], 1:1). WT p53 DBD shows minimal sequence-independent enhanced aggregation at high protein:DNA ratios, less apparent effect at intermediate ratios, and significant a sequence-dependent decrease in aggregation at equimolar ratios. Contact mutants R248Q and R273H maintain the aggregation-enhancing effect at high ratios but loses the inhibitory effect at equimolar concentrations. In contrast, structural mutant R175H shows minimal response to DNA across all concentration ratios, suggesting a fundamental change in DNA interaction. The aggregation pathway (center) progresses from folded monomers through misfolding, pre-fibrillar oligomers, and ultimately to fibril formation. Blue circles represent folded p53 DBD monomers.

### Differential Response of Mutant p53 DBD to DNA Interventions

In contrast to WT p53 DBD, cancer-associated mutants displayed distinct patterns of response to DNA interventions that varied based on both the specific mutation and experimental conditions. The R175H mutant, which exhibited the fastest aggregation kinetics, showed minimal response to most DNA interventions, though interestingly demonstrated increased aggregation across intermediate protein:DNA ratios and with PolyGC sequences in optical density measurements. This complex behavior suggests that even highly destabilized mutants might still engage with DNA in ways that influence their aggregation, though the underlying mechanisms remain to be fully elucidated. R248Q and R273H also showed distinct behaviors, including increased aggregation under various DNA conditions, though through mechanisms that may differ from those seen with WT protein. These mutation-specific responses align with previous studies showing that different p53 mutations require tailored therapeutic approaches. For instance, successful strategies have been developed targeting the Y220X structural mutant through small molecule stabilization (47), while DNA aptamers have shown promise specifically for R175H aggregation (73–75).

### Methodological Considerations and Technical Limitations

Our multi-technique approach provided complementary insights into p53 DBD aggregation dynamics. While AmyTracker630 fluorescence revealed amyloid-like structure formation, optical density measurements captured overall aggregation extent independent of specific structural features, and tryptophan fluorescence monitored local conformational changes. This combination enabled robust validation while revealing distinct aspects of the aggregation process. The utilization of AmyTracker630 as an alternative to ThT proved particularly valuable for monitoring p53 aggregation in the presence of DNA. However, several experimental limitations warrant consideration. The relatively small sample sizes (n=3-5 per condition) may have masked subtle effects, particularly given the inherent variability and stochasticity in protein aggregate formation and kinetics. The statistical analysis, which relied on multiple one-way ANOVAs grouped by concentration followed by post-hoc Dunnett’s tests compared to untreated controls, represents just one analytical approach; alternative statistical methods could yield different significance patterns. While p21-RE effects emerged as the most robust, the absence of false discovery rate corrections likely resulted in more generous significance determinations than a more stringent analysis would support. Though qualitative trends remain apparent, additional replicates would have enhanced statistical power. Moreover, while optical density measurements provided insight into aggregation behavior, their low sensitivity, coupled with the subtle effects observed in mutant:DNA interactions, limited our ability to draw more definitive conclusions. Additionally, our focus on the DBD, while mechanistically informative, does not capture potential contributions from the N-terminal transactivation domains and C-terminal domain in DNA recognition, binding, and overall aggregation (42, 76, 77). Lastly, additional kinetics would provide further insight into possible modulation of the aggregation rate absent in several measurements as end-point analyses except for AT630.

### Biological and Therapeutic Implications

The observed DNA-dependent inhibition of WT p53 DBD aggregation may help explain why WT p53, despite its aggregation-prone nature in vitro, rarely forms aggregates in cells. The ability of WT p53 to bind high-affinity response elements like p21-RE could serve as an endogenous regulatory mechanism that is disrupted by cancer-associated mutations. While our solution biophysics findings illuminate aspects of DNA-mediated effects on p53 DBD aggregation, they also highlight the complexity of these interactions. The observed concentration-dependent responses and mutation-specific behaviors suggest that the presence of DNA may influence p53 stability through multiple mechanisms in addition to or distinct from direct binding, potentially redirecting protein-protein interactions through alternative interfaces. These insights contribute to our understanding of p53 structural dynamics in cellular environments, where various molecular interactions may influence protein behavior. Our findings suggest limited therapeutic potential for exogenously delivered native DNA response elements in modulating mutant p53, emphasizing that specific somatic mutations promote aggregation at the expense of DNA-binding, directly leading to a loss of p53 solubility through biochemical interactions.

Therefore, potent therapeutic strategies targeting p53 aggregation should specifically aim to restore functional binding between oncogenic p53 mutants and endogenous regulatory DNA. This could be achieved by developing modified nucleic acid sequences, such as specific aptamers, optimized for robust binding to mutant p53 and counteracting the observed evasion of DNA-mediated regulation. Alternatively, such modified sequences could function as novel exogenous inhibitors, directly preventing aggregation through distinct mechanisms. To these ends, research has explored various intervention strategies, including aptamers targeting R175H (75), peptide-based approaches (46), and peptide mimetics (48) that have established diverse methods for addressing p53 structural stability. Our work specifically highlights the potential of nucleic acid-based strategies and underscores that continued exploration of these and other approaches is vital. Furthermore, detailed structural characterization of the distinct aggregates formed by WT p53 and different mutants would provide crucial insights into mutation-specific therapeutic strategies.

## Conclusion

In summary, our research demonstrates that high-affinity DNA response elements, such as the p21-RE, are sufficient for regulating the aggregation of WT p53 *in solution*, indicating a possible endogenous mechanism to counter pathological aggregation. However, key somatic mutations implicated in lethal cancers fundamentally alter p53’s interaction with DNA, promoting aggregation and evading the regulatory influence of DNA. These mutation-specific aggregation behaviors highlight the need for tailored therapeutic strategies that restore functional protein-DNA interactions or target the unique pathology of each p53 mutant to combat cancer progression and lethality.

## Materials and Methods

### Protein Expression and Isolation

p53 DBDs comprising residues 94 to 312 were expressed from plasmids containing a 6X His tag/TEV cleavage site with either the WT sequence or specified mutations (R248Q, R273H, R175H) (Genscript, listed in Table S4). BL21(DE3) competent cells (Thermo Fisher Scientific, #EC0114) were transformed and grown overnight at 37°C on LB agar plates containing 40 μg/mL kanamycin (Teknova,#L1053). Starter cultures were prepared in 25 g/L LB media (Millipore Sigma, #L3522) supplemented with 10 μM ZnCl₂ (Rigaku Reagents, #1008318) and 50 μg/mL kanamycin, and grown overnight at 37°C with shaking at 220 RPM. The starter culture was used to inoculate 500 mL LB media containing the same supplements. Cultures were grown at 37°C until reaching OD600 0.7-0.8, induced with 0.5 mM IPTG (Thermo Fisher Scientific, # 15529019), and incubated overnight at 20°C with shaking. All subsequent purification steps were performed at 4°C unless otherwise specified.

Cells were harvested by centrifugation at 6,000 × g for 20 min. Cell pellets were resuspended in 40 mL chilled lysis buffer (1X TBS [Biorad, #1706435], 10% glycerol [Millipore Sigma, #G9012], pH 8.0) containing protease-phosphatase inhibitors (Thermo Fisher Scientific, # A32959). Cells were lysed by sonication using a Qsonica probe sonicator (15 min total, 5 sec on/10 sec off, 60% amplitude). The lysate was clarified by centrifugation at 40,000 × g for 20 min.

The clarified lysate was applied to HisPur™ Ni-NTA Resin (Thermo Fisher Scientific, #88221) pre-equilibrated with 1X TBS (pH 7.5). The column was capped and rotated for 1 h at 4°C. The column was washed with 1X TBS containing 1 mM DTT (GoldBio, #DTT10) until baseline A280 (approximately 40 column volumes [CVs]) monitored using a BioRad Em1 Econo UV Monitor. A subsequent wash was performed with 40 mM imidazole in TBS + 1 mM DTT, collecting fractions up to 20 CVs. Protein was eluted with 500 mM imidazole using the following procedure: 2 mL flush, 6 mL incubation for 3 min, followed by 2 mL without incubation.

Protein purity was assessed by SDS-PAGE before overnight dialysis at 4°C using 3.5-10 kDa MWCO Slide-A-Lyzer™ G3 dialysis cassettes (Thermo Fisher Scientific, #A52967). Dialysis buffer contained 50 mM Tris-HCl (Thermo Fisher Scientific, #J22638-K2), 150 mM NaCl (Sigma, #S6546), 2 mM DTT, pH 8.0, and 3U/mL His-tagged AcTEV protease (Thermo Fisher Scientific, # 12575015). The dialyzed protein was applied to a fresh Ni-NTA column equilibrated with 1X TBS, 10 mM imidazole, 1 mM TCEP (Gold Bio, #TCEP), and 10% glycerol, pH 7.5. Protein was eluted using an imidazole gradient (10-500 mM). Fractions were analyzed by SDS-PAGE, aliquoted, and snap-frozen in liquid nitrogen for storage at -80°C.

### Protein Purification

Frozen protein samples were rapidly thawed and centrifuged at 21,000 × g for 10 min at 4°C. The supernatant was applied to a Capto™ HiRes S 5/50 ion exchange column (Cytiva, #29275877) using an AKTA 25 FPLC system. Samples were loaded at 0.3 mL/min in 20 mM HEPES (Thermo Fisher Scientific, #J60712.AK), pH 7.2. Proteins were eluted using a linear gradient from 0-1 M NaCl in 20 mM HEPES while monitoring A280. Peak fractions were pooled and buffer-exchanged into 1X PBS (Thermo Fisher Scientific, #J75889.K8), 350 mM NaCl, 1 mM TCEP, 10% glycerol, pH 7.4, then aliquoted and snap-frozen.

For final purification, samples were thawed, centrifuged as above, and applied to a Superdex 75 Increase 10/300 GL column (Cytiva, # 29148721) equilibrated in 1X PBS, 1 mM TCEP, pH 7.4. Size exclusion chromatography was performed at 0.7 mL/min, collecting peak fractions for immediate use in downstream experiments. Protein purity was assessed by SDS-PAGE using 4-12% Bis-Tris gels (Thermo Fisher Scientific, #NP0321BOX) in 1X MES buffer (Thermo Fisher Scientific, #NP0002) at 200V for 45 min. Additionally, protein purified by Genscript from the same expression plasmids was used for R175H in thermal denaturation studies, and WT in microscopy. Genscript purified protein underwent the final size exclusion chromatography step described here before experimental use.

### DNA Preparation

DNA sequences were selected based on previous studies: p21-RE and Bax (39) and Poly(GC) (51). A p21-scramble sequence (‘5-AGTACCGTCCGCAGCGACAACTGTCT-3’) was designed using a custom Python script to generate a randomized sequence lacking the consensus elements ’PuPuPuC(A/T)(T/A)GPyPyPy,’ where Pu and Py represent purines and pyrimidines, respectively. HPLC-purified and lyophilized single-stranded DNA oligonucleotides were obtained from Genscript (sequences listed in Table S5). Each oligonucleotide was resuspended in 50 mM Tris-HCl, 137 mM NaCl, pH 8.1, to 200-300 μM final concentration. Complementary strands were annealed by mixing equimolar amounts, heating to 95°C for 2 min in a water-filled heat block followed by gradual cooling to room temperature. Annealing efficiency was assessed using 20% Novex™ TBE PAGE Gels (Thermo Fisher Scientific, #EC63155BOX), with typical yields of 80-95% double-stranded DNA.

Annealed oligonucleotides were snap-frozen and stored at -20°C. Before experiments, samples were rapidly thawed, and concentrations were verified using a NanoDrop™ One Microvolume UV-Vis Spectrophotometer (Thermo Fisher Scientific). Working stocks were prepared in DNA Lo-Bind tubes (Eppendorf, #022431021) at 40X final concentration in 50 mM Tris-HCl, 137 mM NaCl, pH 8.1, maintained on ice throughout sample preparation.

### Sample Preparation for Plate Based Assays

All plate reader assays used a standardized sample preparation protocol. DNA stocks (40X) were prepared in DNA Lo-Bind tubes on ice using 50 mM Tris-HCl, 137 mM NaCl, pH 8.1. Final samples were assembled in 1.5 mL low protein binding tubes (Thermo Fisher Scientific, #90410) by sequential addition of: buffer (1X PBS, 1 mM TCEP, pH 7.4), aggregate dye (ThT or AmyTracker), DNA, and finally 5 μM protein immediately before plate loading. Samples were mixed by gentle inversion (10 times), pulse centrifuged, and 50 μL was transferred in triplicate to microplates using E1-ClipTip™ Bluetooth™ Electronic Multichannel Pipettes (Thermo Fisher Scientific). Samples were added directly to well bottoms, with bubbles removed before sealing. When necessary, plates were centrifuged at 300 RCF for 2 min at 4°C. All plates were sealed with Microseal ’B’ PCR Plate Sealing Film (Bio-Rad, #MSB1001), except for assays requiring endpoint top reads.

### ThT Fluorescence

ThT assays were performed in 96-well Half Area Black/Clear Flat Bottom Polystyrene Non-Binding Surface Microplates (Corning, #3881) using a Molecular Devices M3 plate reader maintained at 37°C. ThT powder (MilliporeSigma, #596200) was stored in a low-humidity desiccator at room temperature and reconstituted in water to 40X stock concentration. ThT concentration was verified using the extinction coefficient ε412 = 31,600 M⁻¹ cm⁻¹ (78), filtered through 0.22 μm filters, and stored at 4°C in the dark for up to one week. Final ThT concentration in samples was 20 μM.

Fluorescence measurements were acquired using bottom reads with medium PMT setting and ten flashes per read. Data was collected every 4 minutes for 24 hours using excitation/emission wavelengths of 440/480 nm and a 475 nm auto cutoff.

### AmyTracker Fluorescence

AmyTracker630 (Ebba Biotech, 1 mg/mL aqueous solution) was diluted to approximately 40X working concentration in 1X PBS, 1 mM TCEP, pH 7.4, immediately before use, based on the molecular weight of 660 ± 30 g/mol (79). Measurements were performed using the same microplate format as ThT assays. Optimal excitation/emission wavelengths were determined to be 490/606 nm with a 590 nm cutoff. Data was collected every 10 minutes for 24 hours; longer intervals were used to minimize mechanical perturbation during the saturation phase.

### Internal Tryptophan Fluorescence

Intrinsic fluorescence measurements were performed in 384-well Low Volume Black Flat Bottom Polystyrene Non-Binding Surface Microplates (Corning, #3820) with 40 μL sample volume. Initial measurements were taken at room temperature using a Perkin Elmer EnVision® 2105 multimode plate reader with monochromator-based top reads. Single-point measurements used excitation/emission of 294/365 nm, while emission scans were performed with 294 nm excitation, scanning 300-500 nm in 5 nm steps.

After initial readings, plates were sealed and incubated at 37°C without shaking for 24 hours. Following incubation, plates were cooled to room temperature and unsealed before final measurements.

### Optical Density Measurements

Optical Density was assessed as an endpoint measurement for pre- and post-aggregation samples using 96-well Half Area Black/Clear Flat Bottom Polystyrene Non-Binding Surface Microplates (Corning, #3881). Absorbance was measured at 350 nm and 400 nm using the multipoint scan setting on a Molecular Devices M3 plate reader. Measurements were taken before and after 24- hour incubation at 37°C.

### Microscopy

All imaging was performed using an ImageXpress Confocal HT.ai High-Content Imaging System (Molecular Devices) equipped with a 40X/1.15 NA CFI APO LWD Lambda S water immersion objective (Nikon, #MRD77410) and a 60 μm pinhole. Time-course imaging was conducted at 37°C, capturing single-site images at the center of each well at 10-minute intervals. Samples were prepared in 96-well Half Area Black/Clear Flat Bottom Polystyrene Non-Binding Surface Microplates sealed with Microseal ’B’ PCR Plate Sealing Film.

### Data Analysis

Data analysis was performed using Python 3.12.2 and Visual Studio Code 1.98.0 for custom model fitting and visualization, alongside GraphPad Prism 10.4.2 for further visualization and statistical analyses. InCarta 2.1 (Molecular Devices) quantified microscopy data, while ImageJ 2.9.0/1.54f handled general image processing. Visualization of PDB: 1TUP was facilitated by PyMOL version 3.1.3.1. Scientific illustrations were created using BioRender, and Adobe Illustrator version 29.2.1 was used for figure creation. For spectroscopic measurements (ThT, AmyTracker, and Optical Density), background buffer signal was subtracted, and aggregation kinetics were fitted to a one-phase association equation:

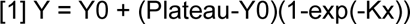

Here, Y0 represents the initial fluorescence, Plateau indicates the saturation fluorescence, K denotes the rate constant (min −1), and x signifies time in minutes. The aggregation half-times (t1/2 =ln(2)/K) were obtained from this equation, with longer half-times suggesting slower aggregation. For spectroscopic techniques, statistical significance was assessed using one-way ANOVA categorized by concentration, followed by Dunnett’s post-hoc tests to compare the conditions against untreated controls (with a threshold of alpha < 0.05). Microscopy data were evaluated using a two-tailed t-test (alpha < 0.05). Raw data and source code available upon request.

### Use of Artificial Intelligence

Artificial intelligence tools such as Grammarly, ChatGPT 4o, Claude-3-5-Sonnet, and Gemini 2.5 Pro were used for editing purposes in the writing of this manuscript. Example prompt: “Please assess the structure and flow of section ‘X’ and identify improvements, inaccuracies, or missing details.” Additionally, GitHub Copilot was used in Python script development and annotation.

**Table 1.** Cancer Type Abbreviations.

| **Cancer Type** | **Abbreviation** |
| --- | --- |
| *Adrenocortical Carcinoma* | ACC |
| *Leukemia* | LEUK |
| *Bladder Cancer* | BLCA |
| *Mature B-Cell Neoplasms* | BNEOP |
| *Breast Cancer* | BRCA |
| *Melanoma* | MEL |
| *Cervical Cancer* | CESC |
| *Non-Small Cell Lung Cancer* | NSCLC |
| *Cholangiocarcinoma* | CHOL |
| *Ovarian Epithelial Tumor* | OET |
| *Colorectal Cancer* | CRC |
| *Pancreatic Cancer* | PAAD |
| *Endometrial Cancer* | EC |
| *Pleural Mesothelioma* | MESO |
| *Esophagogastric Cancer* | EGAC |
| *Prostate Cancer* | PC |
| *Glioblastoma* | GBM |
| *Renal Clear Cell Carcinoma* | ccRCC |
| *Glioma* | GLIOMA |
| *Renal Non-Clear Cell Carcinoma* | nccRCC |
| *Head and Neck Cancer* | HNC |
| *Sarcoma* | SARC |
| *Hepatobiliary Cancer* | HBC |
| *Thymic Epithelial Tumor* | TET |

## Acknowledgments

I extend my gratitude to the Levine Lab, along with Profs. Andrew Miranker and Karla Neugebauer at Yale University for their contributions to this project.

