## Supplemental Figures and Tables for "Consensus DNA Inhibits p53 Aggregation but Fails to Rescue Mutants"

### Supporting Information

#### SI Methods

##### Gel Shift Assays

Gel shift assays were prepared in 20 µL total volume containing 5 µL of protein:DNA mixture (9:1 ratio), 2 µL of Novex™ Hi-Density TBE Sample Buffer (Thermo Fisher Scientific, #LC6678), and non-DEPC treated water (Thermo Fisher Scientific, #AM9939). For 1:1 ratio samples, the mixture was diluted 10-fold, while 454:1 ratio samples were prepared at 2X concentration. Samples were gently mixed by inversion, pulse centrifuged and maintained on ice until loading.

Samples (10 µL/well) were analyzed on 6% DNA retardation gels (Thermo Fisher Scientific, #EC6365BOX) using an XCell SureLock Gel Tank (Thermo Fisher Scientific). Electrophoresis was performed at room temperature for 50 minutes at 150V in 0.5X TBE buffer. Gels were rinsed with 0.5X TBE buffer and stained with 1X SYBR™ Gold Nucleic Acid Gel Stain (Thermo Fisher Scientific, #S11494) for 20 minutes at room temperature with gentle rocking. Imaging was performed using a ChemiDoc MP system (Bio-Rad) with SYBR Gold channel settings as specified in the results section.

##### Thermal Melt Assay

Thermal stability measurements were conducted in quadruplicate using a QuantStudio 7 real-time PCR system (Applied Biosystems) in 384-well plates. Samples were prepared using the Protein Thermal Shift™ Dye Kit (Applied Biosystems, #4461146) according to manufacturer's protocol, with final protein concentration of 1 µg/µL in 20 µL total volume. Temperature was ramped from 15°C to 99°C at 0.1335°/sec. Melting temperatures (Tm) were determined from the derivative of the melt curve using Protein Thermal Shift software (Applied Biosystems). Raw data and analysis were exported and plotted using GraphPad Prism 10. Thermal melts were performed on both in-house and Genscript-purified WT, R248Q, and R273H proteins, while R175H analysis used only Genscript-purified protein.

##### Internal Tryptophan Fluorescence

To minimize inner filter effects from DNA, excitation wavelength was optimized to 294 nm, corresponding to ~0.1 absorbance units for DNA oligos. While 5 µM samples showed significant signal decrease across all oligos, consistent initial intensity values across DNA concentrations (Fig. 3D) suggest minimal inner filter effects.

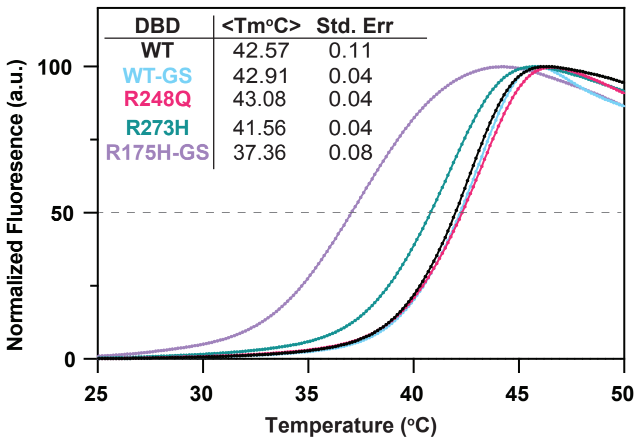


***S1. Thermal stability analysis confirms expected destabilization of R273H and R175H p53 DBD mutants compared to wild-type protein.*** *Normalized fluorescence thermal melt curves for wild-type p53 DBD (WT), Genscript-sourced wild-type (WT-GS), and mutants R248Q, R273H, and R175H-GS (n = 4). The grey dashed line indicates 50% normalized fluorescence, corresponding to the melting temperature (Tm) where half the protein population is predicted to be unfolded based on SYPRO Orange binding. Data points represent mean values with standard error shown. Consistent with previous literature, R273H and R175H demonstrate reduced thermal stability compared to wild-type protein.*

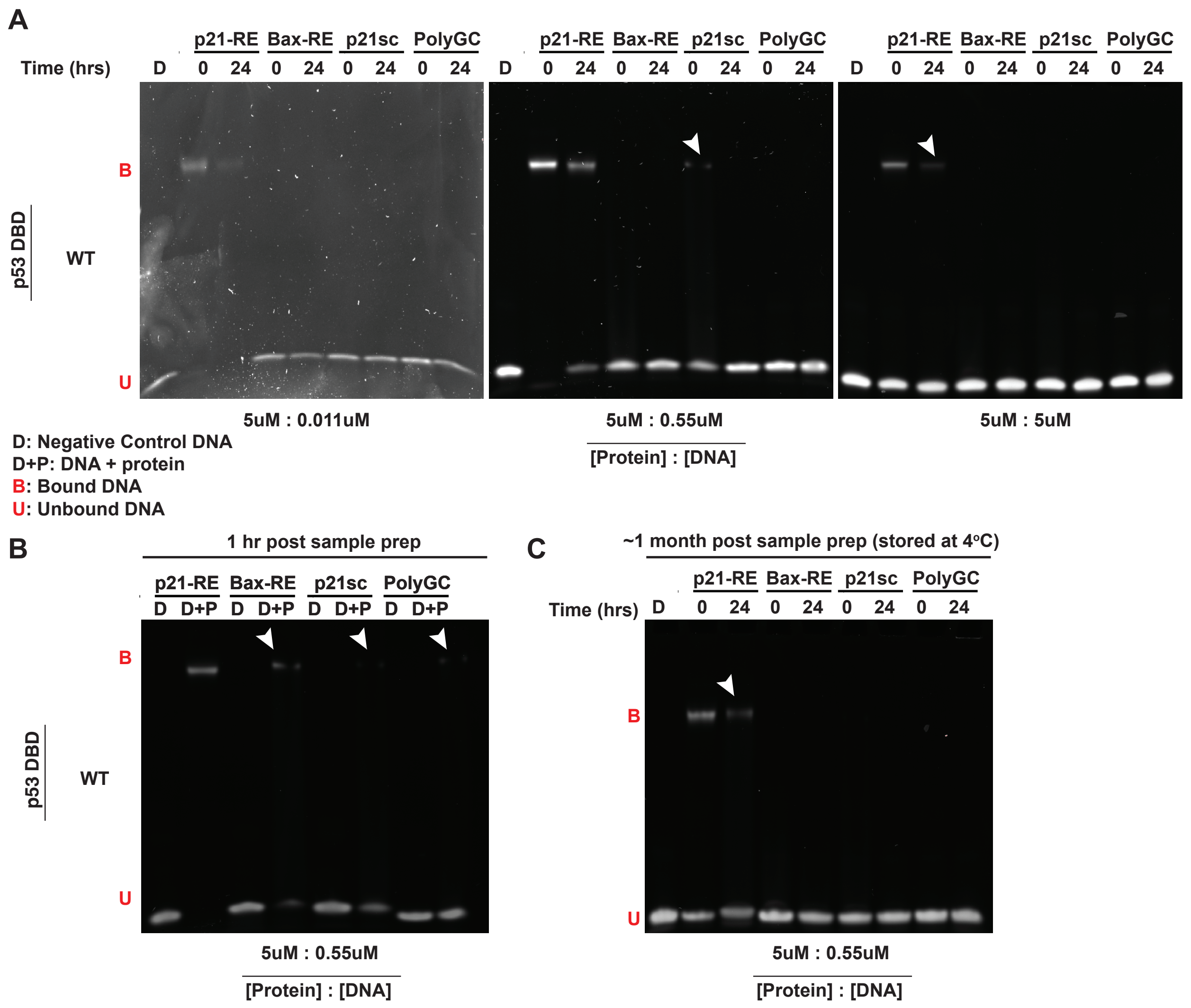


***S2. Gel shift analysis demonstrates time- and concentration-dependent DNA binding patterns of p53 DBD.*** *A) Gel shift analysis of p53 DBD-DNA interactions at protein:DNA ratios of 1:1, 9:1, and 454:1. After sample preparation, aliquots were either maintained at 4°C (0 hr samples) or incubated at 37°C for 24 hrs to stimulate aggregation, then analyzed simultaneously. DNA sequences tested include p21-RE, Bax-RE, p21sc, and polyGC. B) 9:1 ratio of protein to DNA following approximately 1hr of incubation at 4°C. C) Samples were prepared as in A, and stored at 4°C for approximately 1 month. All samples were resolved on 6% retardation gels and visualized using SYBR Gold staining. White arrows indicate faint bands corresponding to protein-DNA complexes.*

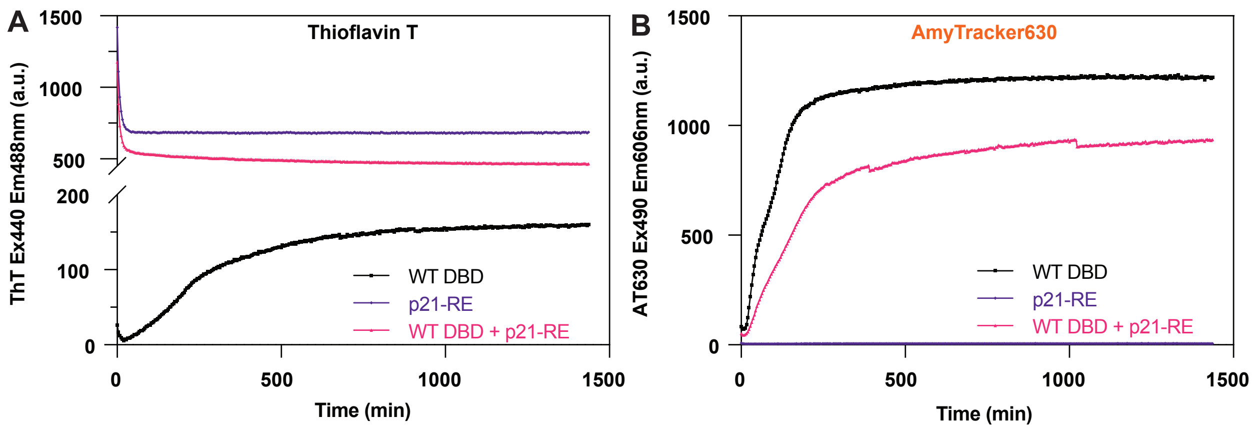


***S3. Comparison of ThT and AmyTracker630 for monitoring p53 DBD aggregation in the presence of DNA.*** *A) Thioflavin T (ThT, 20 µM) fluorescence monitoring of wild-type p53 DBD (15 µM) aggregation alone, p21-RE DNA (10 µM) alone, and their combination over time. ThT signal decreases for p21-RE and WT DBD + p21-RE samples while increasing for WT DBD alone. B) AmyTracker630 (1:40 dilution from 1 mg/mL stock) fluorescence shows consistent signal increase over time for all samples, demonstrating improved detection of protein aggregation in the presence of DNA compared to ThT. Data represent mean traces from two independent measurements with background fluorescence subtracted.*

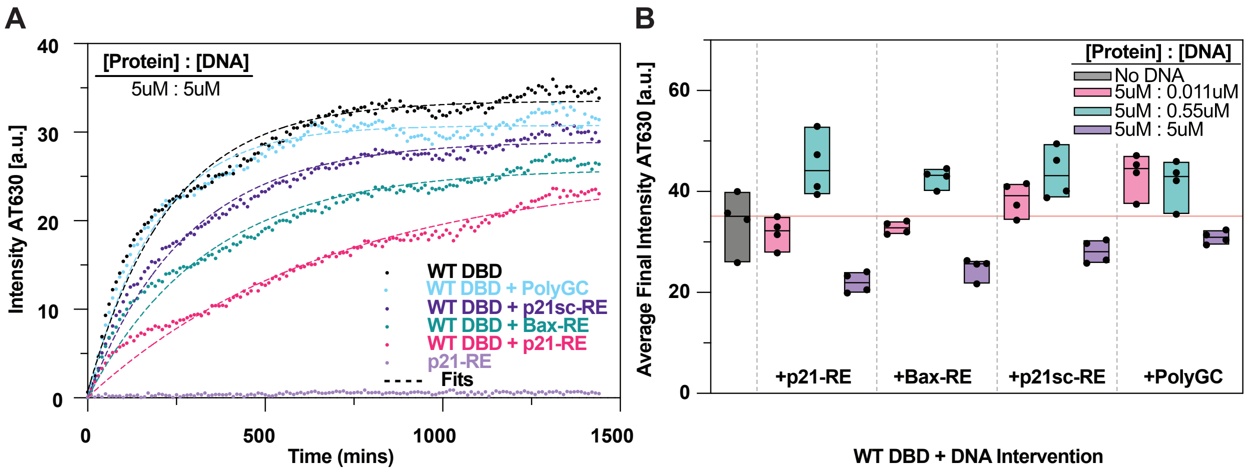


***S4. Absolute AmyTracker630 fluorescence data confirms DNA sequence- and concentration-dependent effects on WT p53 DBD aggregation.*** *A) Representative time course curves and fits of raw (unnormalized) AmyTracker630 intensity for WT p53 DBD over 24 hours with and without consensus DNA at equimolar protein to DNA concentration (5 : 5 μM) (n = 4). B) Final aggregation extent calculated as the average AmyTracker630 intensity over the last six time points for each replicate.*

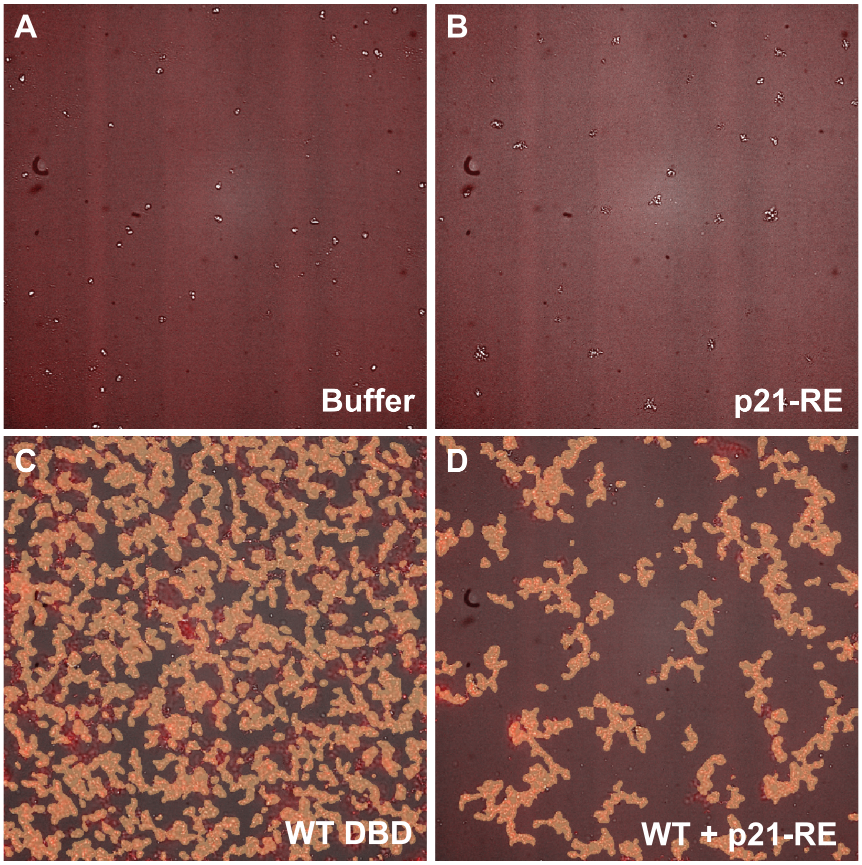


***S5. InCarta SINAP model detection of p53 DBD aggregates.*** *Representative composite images showing transillumination (grey) and AmyTracker630 fluorescence (red) channels with SINAP model-identified aggregates highlighted by orange masks for: (A) Buffer control, (B) p21-RE DNA alone, (C) wild-type p53 DBD alone, and (D) wild-type p53 DBD with p21-RE DNA. The machine learning-based SINAP model was trained to specifically identify protein aggregates in these samples.*

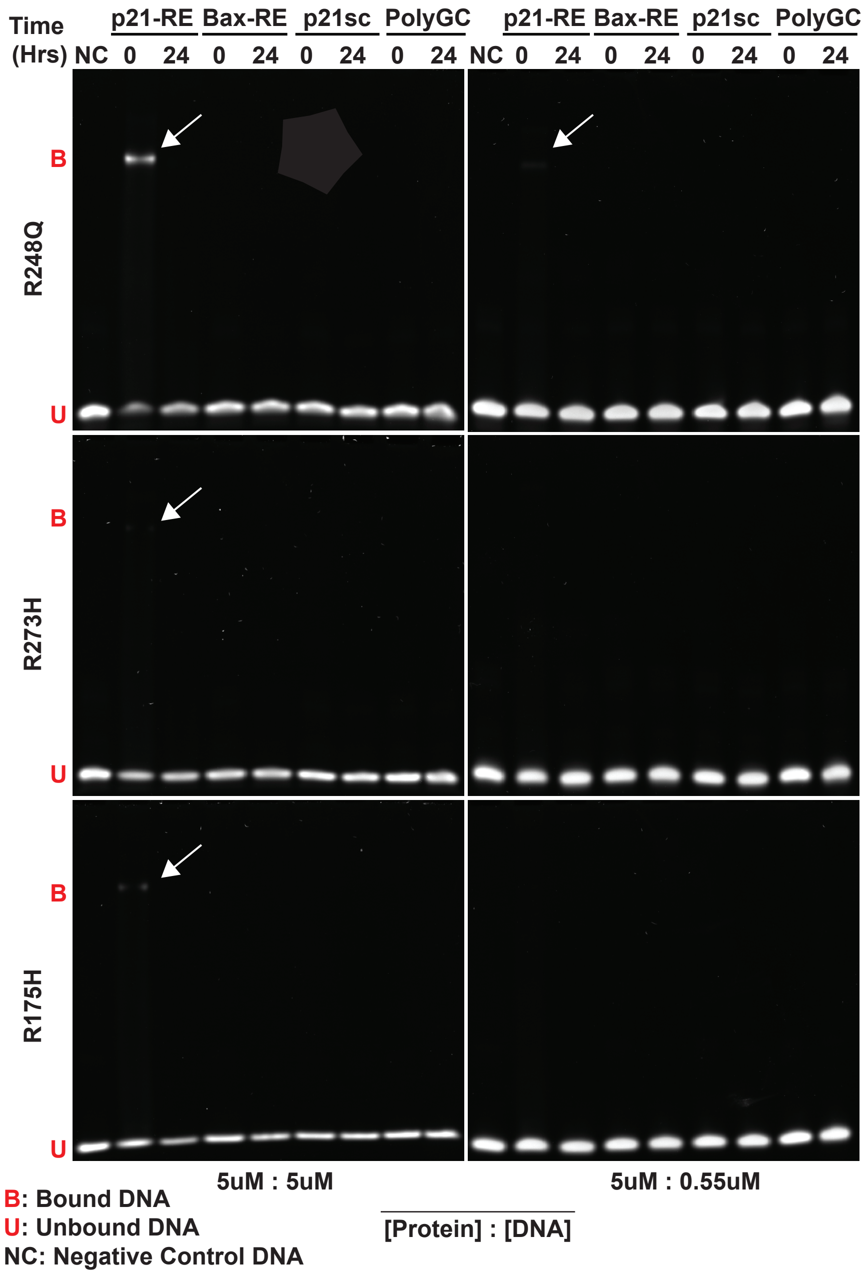


***S6. Gel shift analysis reveals altered DNA binding patterns of p53 mutants.*** *Gel shift analysis of R248Q, R273H, and R175H p53 DBD mutants with DNA at protein:DNA ratios of 1:1 and 9:1. Samples were analyzed at 0 hours (maintained at 4°C) and after 24 hours of aggregation conditions. DNA sequences tested include p21-RE, Bax-RE, p21sc, and polyGC. White arrows indicate faint bands corresponding to protein-DNA complexes at 0 hours. Samples were resolved on 6% retardation gels and visualized using SYBR Gold staining.*

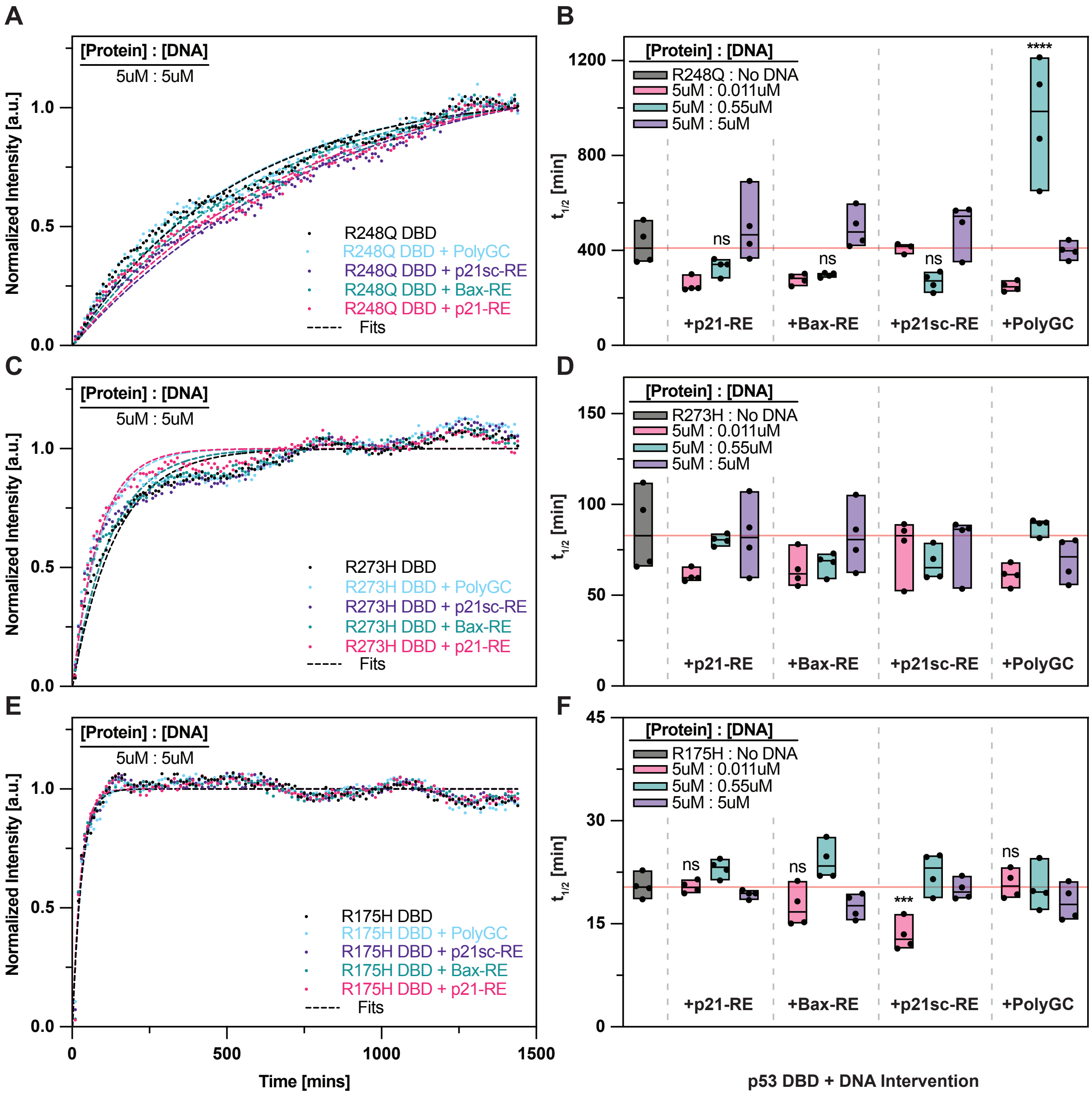


***S7. p53 DBD hotspot mutants exhibit distinct aggregation kinetics with minimal DNA-dependent effects.*** *A) Representative time course curves and fits of normalized AmyTracker630 intensity for R248Q, C) R273H, and E) R175H p53 DBDs over 24 hours ± consensus DNA at equimolar concentrations (5:5 μM) (n = 3-4). B) t-half values for R248Q, D) R273H, and F) R175H p53 DBDs from independent one-phase association fits to each replicate. . Each set of concentrations for each protein was analyzed using a one-way ANOVA followed by a post hoc Dunnett's test. If no symbols are displayed, the sample failed to reject the ANOVA. Dunnett's adjusted P-values were mapped to significance symbols as follows: ns ≥ 0.05 ; ✱ : P ≤ 0.05 ; ✱✱ : P ≤ 0.01 ; ✱✱✱ : P ≤ 0.001 ; ✱✱✱✱ : P ≤ 0.0001.*

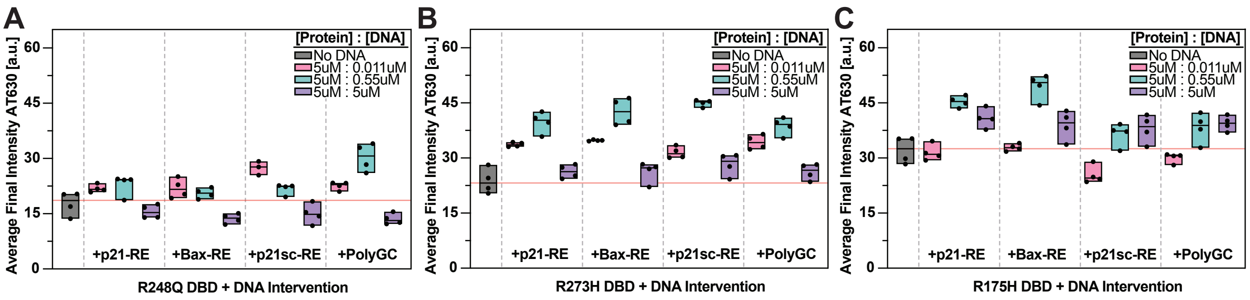


***S8. Mutation-specific differences in final aggregation extent of p53 DBD variants.*** *Final aggregation extent shown as average AmyTracker630 intensity over the last six time points for each replicate of A) R248Q, B) R273H, and C) R175H p53 DBD variants with and without consensus DNA at varying protein-to-DNA ratios.*

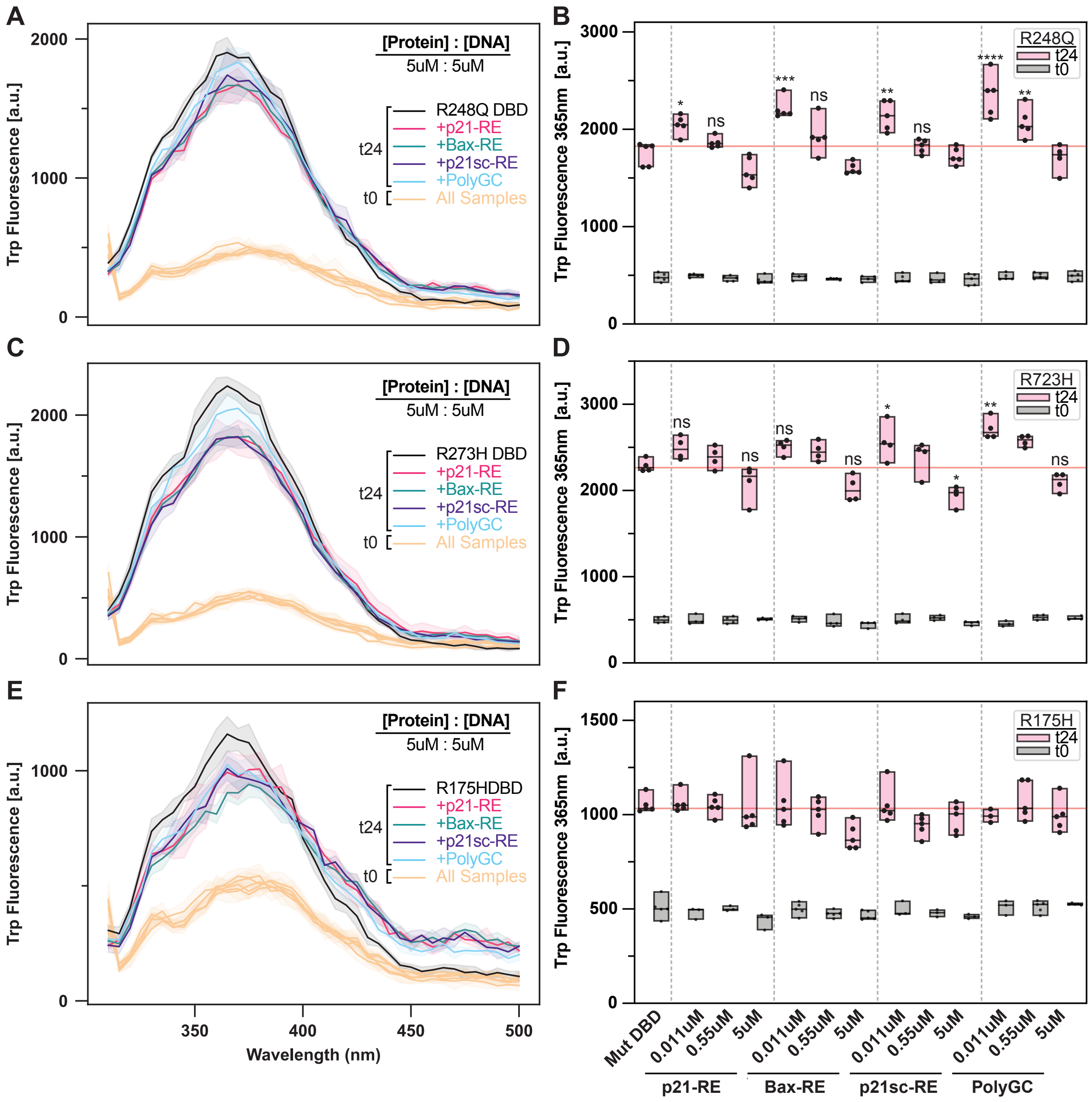


***S9. p53 DBD hotspot mutants show reduced tryptophan fluorescence, with R248Q and R273H maintaining WT-like DNA response patterns.*** *A) Emission Scan for Internal Trp Fluorescence of R248Q, C) R273H, and E) R175H p53 DBDs and boxplots of emission values at 365nm for B) R248Q, D) R273H, and F) R175H (n = 3-5). Internal tryptophan fluorescence was measured in triplicate to quintuplicate before aggregation (0 hours) and after 24 hours of incubation at 37°C to initiate aggregation. Samples were excited at 298nm and scanned from 310 to 500nm. Each set of concentrations for each protein was analyzed using a one-way ANOVA followed by a post hoc Dunnett's test. If no symbols are displayed, the sample failed to reject the ANOVA. Dunnett's adjusted P-values were mapped to significance symbols as follows: ns ≥ 0.05 ; ✱ : P ≤ 0.05 ; ✱✱ : P ≤ 0.01 ; ✱✱✱ : P ≤ 0.001 ; ✱✱✱✱ : P ≤ 0.0001.*

#### *Table S1. AmyTracker630: One-Way ANOVAs and Posthoc Dunnett's P-values*

| Technique | DBD | Sample Group | ANOVA P-Val | ANOVA f-test | ANOVA DF | Dunnett's P-Val [ DBD + p21 ] | Dunnett's P-Val [ DBD + Bax ] | Dunnett's P-Val [ DBD + p21sc ] | Dunnett's P-Val [ DBD + PolyGC ] |
|---|---|---|---|---|---|---|---|---|---|
| AT630 | WT | WT, WT+DNA [0.011uM] | 0.00378487 | 6.191336442 | 4 | 0.006724092 | 0.041214767 | 0.989784468 | 0.013750608 |
| | | WT, WT+DNA [0.55uM] | 4.29E-05 | 14.82309825 | 4 | 0.030443248 | 0.017732935 | 0.01472298 | 0.867949888 |
| | | WT, WT+DNA [5uM] | 1.36E-07 | 36.86881777 | 4 | 4.12E-09 | 0.12274017 | 0.12175312 | 0.921938992 |
| | R248Q | R248Q, R248Q+DNA [0.011uM] | 0.13282235 | 2.090424513 | 4 | x | x | x | x |
| | | R248Q, R248Q+DNA [0.55uM] | 3.31E-06 | 22.61001169 | 4 | 0.654514783 | 0.396757842 | 0.242284432 | 3.15E-05 |
| | | R248Q, R248Q+DNA [5uM] | 0.4442644 | 0.987031615 | 4 | x | x | x | x |
| | R273H | R273H, R273H+DNA [0.011uM] | 0.075104756 | 2.642164571 | 4 | x | x | x | x |
| | | R273H, R273H+DNA [0.55uM] | 0.051456734 | 3.025793308 | 4 | x | x | x | x |
| | | R273H, R273H+DNA [5uM] | 0.758158657 | 0.468341167 | 4 | x | x | x | x |
| | R175H | R175H, R175H+DNA [0.011uM] | 0.000556661 | 9.278967398 | 4 | 0.999959964 | 0.163358726 | 0.000716378 | 0.999427209 |
| | | R175H, R175H+DNA [0.55uM] | 0.179440844 | 1.810012616 | 4 | x | x | x | x |
| | | R175H, R175H+DNA [5uM] | 0.160514978 | 1.913173542 | 4 | x | x | x | x |

#### *Table S2. Trp Fluorescence: One-Way ANOVAs and Posthoc Dunnett's P-values*

| Technique | DBD | Sample Group | ANOVA P-Val | ANOVA f-test | ANOVA DF | Dunnett's P- Val [ DBD + p21 ] | Dunnett's P-Val [ DBD + Bax ] | Dunnett's P-Val [ DBD + p21sc ] | Dunnett's P-Val [ DBD + PolyGC ] |
|---|---|---|---|---|---|---|---|---|---|
| Trp Fluorescence | WT | WT, WT+DNA [0.011uM] | 0.021172038 | 3.992934092 | 4 | 0.708303323 | 0.995312983 | 0.234989623 | 0.224662538 |
| | | WT, WT+DNA [0.55uM] | 0.058352356 | 2.896496933 | 4 | x | x | x | x |
| | | WT, WT+DNA [5uM] | 1.23E-09 | 72.68971345 | 4 | 9.99E-16 | 3.60E-07 | 1.31E-07 | 0.000108833 |
| | R248Q | R248Q, R248Q+DNA [0.011uM] | 4.67E-05 | 11.68546266 | 4 | 0.708303323 | 0.995312983 | 0.234989623 | 0.224662538 |
| | | R248Q, R248Q+DNA [0.55uM] | 0.008581 | 4.590903291 | 4 | 0.020462021 | 0.102237273 | 0.081204263 | 0.275725609 |
| | | R248Q, R248Q+DNA [5uM] | 0.085876686 | 2.382187736 | 4 | x | x | x | x |
| | R273H | R273H, R273H+DNA [0.011uM] | 0.010059144 | 4.88580372 | 4 | 0.17309891 | 0.13450823 | 0.044476213 | 0.002185441 |
| | | R273H, R273H+DNA [0.55uM] | 0.061081223 | 2.849937815 | 4 | x | x | x | x |
| | | R273H, R273H+DNA [5uM] | 0.038804261 | 3.322539791 | 4 | 0.185238723 | 0.055020003 | 0.011608806 | 0.221161822 |
| | R175H | R175H, R175H+DNA [0.011uM] | 0.831387136 | 0.363437839 | 4 | x | x | x | x |
| | | R175H, R175H+DNA [0.55uM] | 0.055092213 | 2.777999628 | 4 | x | x | x | x |
| | | R175H, R175H+DNA [5uM] | 0.085111573 | 2.390057046 | 4 | x | x | x | x |

#### *Table S3. Optical Density: One-Way ANOVAs and Posthoc Dunnett's P-values*

| Technique | DBD | Sample Group | ANOVA P-Val | ANOVA f-test | ANOVA DF | Dunnett's P- Val [ DBD + p21 ] | Dunnett's P-Val [ DBD + Bax ] | Dunnett's P-Val [ DBD + p21sc ] | Dunnett's P-Val [ DBD + PolyGC ] |
|---|---|---|---|---|---|---|---|---|---|
| Optical Density | WT | WT, WT+DNA [0.011uM] | 1.50E-09 | 70.72916611 | 4 | 0.829268472 | 1.83E-09 | 5.46E-09 | 2.87E-07 |
| | | WT, WT+DNA [0.55uM] | 0.011948351 | 4.672019077 | 4 | 0.006313139 | 0.994379112 | 0.165806378 | 0.619063419 |
| | | WT, WT+DNA [5uM] | 3.91E-08 | 49.92171495 | 4 | 4.69E-05 | 2.57E-05 | 0.000992269 | 0.093420363 |
| | R248Q | R248Q, R248Q+DNA [0.011uM] | 3.65E-05 | 21.57887915 | 4 | 0.552338079 | 0.060398358 | 0.021808852 | 2.82E-05 |
| | | R248Q, R248Q+DNA [0.55uM] | 0.002478157 | 8.270521559 | 4 | 0.000698919 | 0.681120113 | 0.614261771 | 0.500761549 |
| | | R248Q, R248Q+DNA [5uM] | 1.49E-06 | 25.62418826 | 4 | 0.001606391 | 0.95313122 | 0.712870531 | 0.000223046 |
| | R273H | R273H, R273H+DNA [0.011uM] | 0.012751597 | 4.592377353 | 4 | 0.037812293 | 0.011562883 | 0.012234438 | 0.484839921 |
| | | R273H, R273H+DNA [0.55uM] | 0.000745047 | 8.758492296 | 4 | 0.40099053 | 0.99752548 | 0.024250494 | 0.000532741 |
| | | R273H, R273H+DNA [5uM] | 1.43E-05 | 17.84085969 | 4 | 0.000202792 | 1.87E-06 | 0.000204598 | 0.045235007 |
| | R175H | R175H, R175H+DNA [0.011uM] | 0.031944925 | 3.532650583 | 4 | 0.211437389 | 0.961205022 | 0.74158105 | 0.014400966 |
| | | R175H, R175H+DNA [0.55uM] | 3.77E-05 | 15.15418864 | 4 | 0.000190613 | 0.00015916 | 8.94E-06 | 0.000323674 |
| | | R175H, R175H+DNA [5uM] | 2.31E-06 | 23.92307138 | 4 | 0.311939337 | 0.049848682 | 0.93784746 | 1.91E-07 |

#### *Table S4. Genscript Plasmid Sequences*

Construct features: NdeI--ATG--His tag--TEV protease site--**p53**--Stop codon--HindIII

| Construct | Plasmid Backbone | Plasmid Size | Antibiotic Resistance | Copy Number | Quality Grade |
|---|---|---|---|---|---|
| WT | pET-30a(+) | 5973 | Kanamycin | Low | HT Transfection Grade (90%±10% Supercoiled, ≤ 0.1 EU/μg endotoxin) |
| R248Q | pET-30a(+) | 5973 | Kanamycin | Low | HT Transfection Grade (90%±10% Supercoiled, ≤ 0.1 EU/μg endotoxin) |
| R175H | pET-30a(+) | 5973 | Kanamycin | Low | HT Transfection Grade (90%±10% Supercoiled, ≤ 0.1 EU/μg endotoxin) |
| R273H | pET-30a(+) | 5973 | Kanamycin | Low | HT Transfection Grade (90%±10% Supercoiled, ≤ 0.1 EU/μg endotoxin) |

#### *Table S5. DNA Sequences*

| Name | Strand | Sequence 5'->3' | # Bases | Source | Purification |
|---|---|---|---|---|---|
| p21 | Sense | CGCGAACATGTCCCAACATGTTGCGC | 26 | Genscript | HPLC |
| p21 | Antisense | GCGCAACATGTTGGGACATGTTCGCG | 26 | Genscript | HPLC |
| Bax | Sense | CGCTCACAAGTTAGAGACAAGCTTCGC | 27 | Genscript | HPLC |
| Bax | Antisense | GCGAAGCTTGTCTCTAACTTGTGAGCG | 27 | Genscript | HPLC |
| p21-scramble | Sense | AGTACCGTCCGCAGCGACAACTGTCT | 26 | Genscript | HPLC |
| p21-scramble | Antisense | AGACAGTTGTCGCTGCGGACGGTACT | 26 | Genscript | HPLC |
| Poly(GC) | Sense | ATAATTGCGCGCGCGCGCAGGAAA | 24 | Genscript | HPLC |
| Poly(GC) | Antisense | TTTCCTGCGCGCGCGCGCAATTAT | 24 | Genscript | HPLC |
